# The role of secondary structures of peptide polymers on antimicrobial efficacy and antibiotic potentiation

**DOI:** 10.1101/2024.11.19.623429

**Authors:** Swagatam Barman, Alimi Abiodun, Md Waliullah Hossain, Adam Parris, Aiswarya Banshidhar Chandrasseril, Ethan A. Older, Jie Li, Alan W. Decho, Chuanbing Tang

## Abstract

The rise of antibiotic resistance, biofilm formation, and dormant bacterial populations poses serious global health threats. Synthetic antimicrobial peptide (AMP) mimics offer promising alternatives, though the impact of secondary structures in polymeric AMP mimics on antimicrobial efficacy is underexplored. This study investigates chirality-controlled α-peptide polymers (D-PP and DL-PP), synthesized via ring-opening polymerization of allylglycine *N*-carboxy anhydrides and post-polymerization modification through thiol-ene click chemistry. D-PP adopts a stable helical structure under biomimetic conditions, whereas DL-PP remains random. This helical structure enhanced D-PP’s antibacterial and antibiotic potentiation activities, amplifying antibiotic efficacy by 2- to 256-fold across various classes—including tetracyclines, ansamycins, fusidanes, macrolides, cephalosporins, and monobactams—against multidrug-resistant Gram-negative pathogens, while maintaining low hemolytic activity and high protease stability. Mechanistic investigations revealed that D-PP exhibited greater membrane interaction. D-PP and antibiotic combinations eradicated dormant bacterial populations and disrupted biofilms with minimal antimicrobial resistance development. This study paves the way for the rational design of polypeptide-based antimicrobial agents, harnessing chirality and secondary structural features to enhance the efficacy of synthetic antimicrobial peptide mimics.

## INTRODUCTION

To address the escalating crisis of antimicrobial resistance (AMR), short-cationic naturally occurring antimicrobial peptides (AMPs) received significant attention owing to their selective interaction with the negatively charged microbial membrane over the zwitterionic mammalian membranes.^1-3^ The role of secondary structures in AMPs is essential for targeting bacterial membranes, with natural AMPs often adopting helical conformations and/or facial amphiphilicity that enhance antimicrobial activity.^4^ ^5,6^ Despite the broad-spectrum antimicrobial efficacy and strong immunity to microbial resistance, their transition from laboratory to clinics is limited because of protease instability, complexity in synthesis, high manufacture cost and unwanted systemic toxicity. These challenges have driven the design of synthetic AMP mimics. Researchers have explored a variety of peptidomimetics such as α-peptides,^7,8^ β-peptides,^9^ γ-AA peptides,^10^ foldamers,^11^ peptoids,^12^ and cationic amphiphiles.^13-16^ These peptidomimetics served their purpose in preventing protease degradation and enhancing antimicrobial efficacy. However, many of the designs required amino acid sequence specificity and often suffered from synthetic complexity including the involvement of solid-state peptide synthesis.

Though extensive efforts are still being attempted to prepare a vast array of diverse synthetic peptidomimetics to enhance efficacy against microbes, the direct impact of secondary structures on antimicrobial activity remained largely underexplored. There have been limited efforts in designing D-, L- or DL-polypeptides,^6,17-19^ where notably, rigorous comparisons between D-stereo-configured polypeptides and racemic DL-polypeptides, particularly regarding chirality’s effect on structure and antimicrobial efficacy, are lacking. D-peptides generally exhibit greater proteolytic stability compared to their L-peptide counterparts.^20^ This difference is primarily because proteases, the enzymes responsible for breaking down peptides, are stereospecific for the L-configuration of amino acids, which are the naturally occurring form in biological systems.^21,22^ D-peptides, consisting of D-amino acids, are resistant to cleavage by most natural proteases since these enzymes are not adapted to recognize or process the D-amino acid configuration effectively.^20^

Here, we design primary ammonium-bearing allylglycine-based D-polypeptide (D-PP) with DL-polypeptide (DL-PP) as a simple model peptide polymer system and show that the chirality of peptide building blocks strongly influences secondary structure formation and antimicrobial potency. D-PP adopts a stable helical structure under biomimicking environments, while racemic DL-PP lacks a defined conformation. Despite this model system is not intended to achieve optimization of structures and compositions, our findings demonstrate that the helical D-PP significantly enhances antibacterial activity compared to the non-helical DL-PP. Additionally, D-PP’s helical conformation also boosts its ability to potentiate antibiotics, particularly those limited by membrane permeability or efflux mechanisms. Mechanistic studies revealed that D-PP’s helical structure promotes stronger interactions with bacterial membranes, explaining its improved performance relative to DL-PP. We also evaluated D-PP-antibiotic combinations against biofilms and dormant bacterial populations, which are major contributors to persistent infections. The results highlight the enhanced therapeutic potential of D-PP, with resistance development studies indicating that D-PP-antibiotic combinations may reduce the emergence of resistant strains.

This study is the first of its kind to compare D- and DL-based peptide polymers, offering key insights into how chirality and secondary structures influence the efficacy of synthetic AMP mimics. These findings open new possibilities for the rational design of polypeptide-based antimicrobial agents that leverage structural advantages to combat AMR and address the growing threat of resistant infections.

## RESULTS AND DISCUSSION

### Synthesis and secondary structures of peptide polymers

Employing ring-opening polymerization (ROP) and post-polymerization modification,^23^ we synthesized D-PP and DL-PP. The molecular design was inspired by the facially amphiphilic architecture of natural and synthetic AMPs,^24,25^ which usually adopt helical structures. The peptide polymers were prepared in three steps, including monomer synthesis, polymerization, and post-polymerization modification (Figure 1a). D and DL stereoisomers of allylglycine were converted to *N*-carboxy anhydride (NCA) derivatives by reacting with triphosgene in the presence of propylene oxide (PO), which acted as a proton scavenger.^26^ Further, ROP of D- or DL-allylglycine NCAs was carried out using benzylamine as an initiator under an inert atmosphere at room temperature for 24 h. ^1^H NMR spectra of poly(D- or DL-allylglycine) show the terminal protons (C*H₂*CH-) and the single proton (CH₂C*H*) of the double bond at the pendant chain, relative to the aromatic protons from the end group (Figures S1 and S2: a and b protons in the red spectrum). While the degree of polymerization (DP) can be facilely tuned by changing molar ratios of monomer to initiator, polymers with DP of approximately 20 were prepared, which is in the typical range of AMPs. Additionally, the lack of both symmetric and asymmetric stretching vibrations (∼1754 and ∼1827 cm^-1^) typically associated with the carbonyl group (non-amide) in the FT-IR spectra provided evidence for the successful polymerization of the cyclic NCA monomers, resulting in the formation of the peptide polymers, poly(D- or DL-allylglycine) (Figures S7 and S8). These precursor polymers were post-functionalized through thiol-ene click reaction between free terminal alkenes and cysteamine hydrochloride. The complete disappearance of the double bond protons at 5.2-5.7 ppm indicated the efficient click reaction, resulting in the formation of amphiphilic water-soluble peptide polymers, D-PP (chiral centre: R configuration) and DL-PP (chiral centre: R/S configuration) (Figures S1 and S2). We employed ^1^H-NMR, ^13^C-NMR, FT-IR and mass spectrometry (supplementary Figures: S1-S8) to fully characterize both the polymers and monomers.

**Figure 1.**
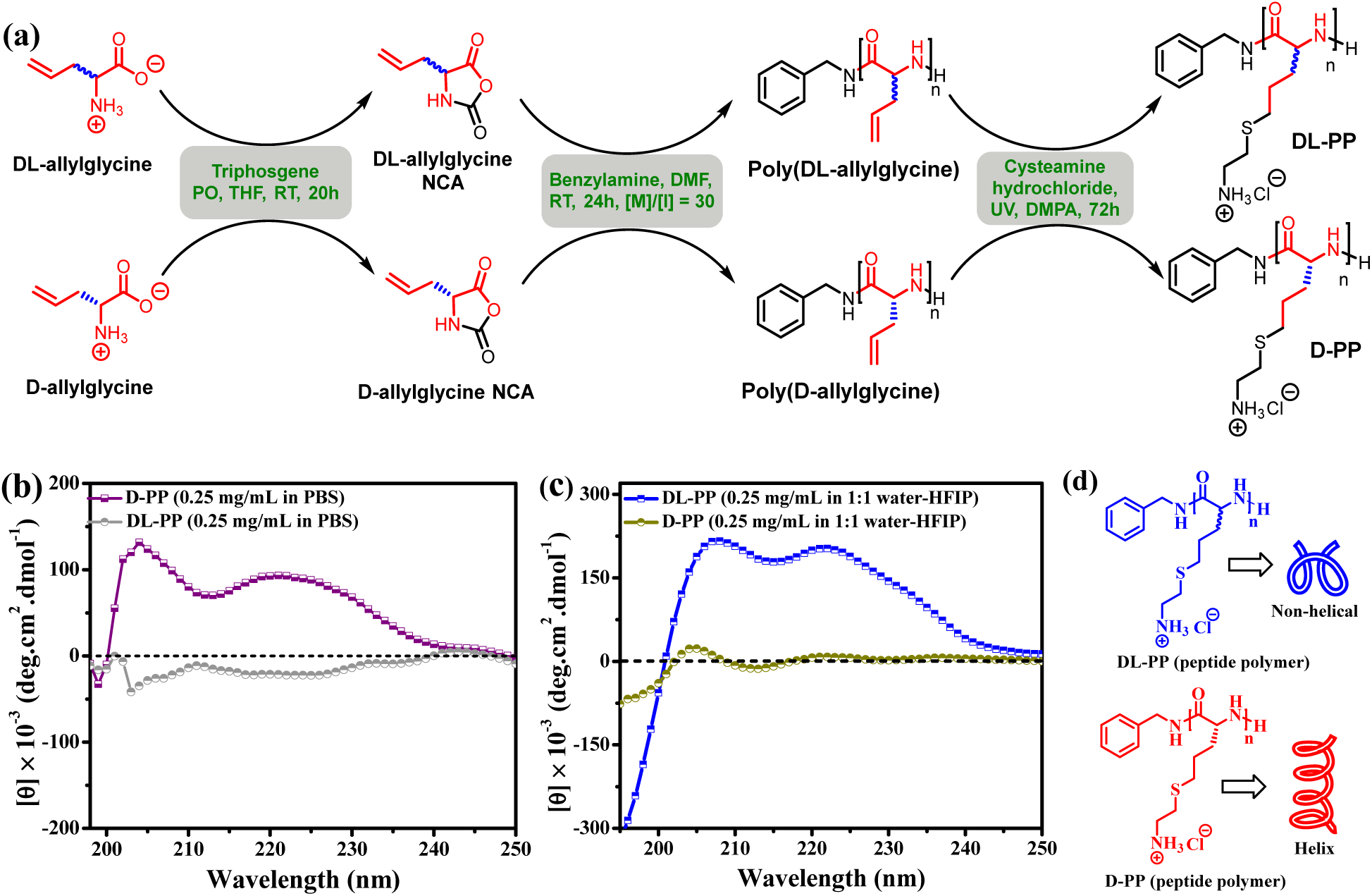
Synthesis and secondary structures of peptide polymers. (a) Ring-opening polymerization of *N*-carboxy anhydride and post-polymerization modification; (b) CD spectra of DL-PP and D-PP in PBS; (c) CD spectra of DL-PP and D-PP in a mixture of HFIP and water (v/v = 1/1); and (d) The schematic representation of DL-PP without a defined secondary structure and D-PP with a helical structure.

The precursor polymers and corresponding D-PP and DL-PP were characterized by circular dichroism (CD) spectroscopy (Figure 1 and Figure S9). A CD spectrum of poly(D-allylglycine) in hexafluoro isopropyl alcohol (HFIP), an amphiphilic solvent known for mimicking the membrane environment in peptide and protein studies, revealed two maxima at 203 nm and 220 nm (Figure S9). This indicated the formation of an α-helix-like secondary structure. In comparison, both maxima are entirely missing for the DL-analogue, indicating a random conformation with the absence of a defined secondary structure. After installing cationic primary ammonium at the pendant groups via the post-polymerization modification of the precursor, the resultant D-PP maintained its helical conformation (double maxima at 208 nm and 222 nm) when dissolved in a mixture of HFIP and water (v/v=1:1) (Figure 1c), while DL-PP still exhibited a lack of defined secondary structures. Both polypeptides were also dissolved in the phosphate buffer saline (PBS) solution. D-PP also displayed a helix, as evidenced by the double maxima at 204 nm and 222 nm in the CD spectrum (Figure 1b). While DL-PP could not adopt a helical conformation in PBS, similar to its corresponding precursor, it is worth mentioning that adopting a helix-like conformation in PBS by the D-PP required a long incubation period (72h) at 37 ^°^C. In contrast, the formation of helix was almost instantaneous in HFIP/water. It has been suggested that a mixture of water and fluorinated alcohol (e.g. HFIP or trifluoro ethanol) mimics the amphiphilic nature of cellular membranes.^27,28^ These results demonstrated that the D-peptide polymer readily adopts a helix structure upon interaction with the biomembrane-mimicking environment (Figure 1d).

### Dual efficacy as an antimicrobial agent and a universal adjuvant

To understand the impact of secondary structures, the antimicrobial activity of both helical (D-PP) and non-helical (DL-PP) peptide polymers was evaluated against various strains of Gram-positive *S. aureus* and Gram-negative bacteria using micro broth dilution assays according to CLSI guidelines.^29^ Both polymers displayed appreciable antimicrobial activity with minimum inhibitory concentrations (MICs) in the range of 4-16 µg/mL against Gram-positive bacteria, including MRSA (Figure 2a). However, MICs were significantly higher (64-256 µg/mL) against MDR Gram-negative bacteria (except for MDR *P. aeruginosa* ATCC BAA 2108, where MIC of D-PP was 16 µg/mL). Notably, D-PP showed 2 to 4-fold lower MICs against both Gram-positive and Gram-negative strains compared to DL-PP (Figure 2b), indicating the helical polymer’s enhanced efficacy towards bacteria. It should be noted that both polymers show very low hemolysis (<10%) against mouse erythrocytes up to tested concentrations of 1024 µg/mL (Figure 2c).

**Figure 2.**
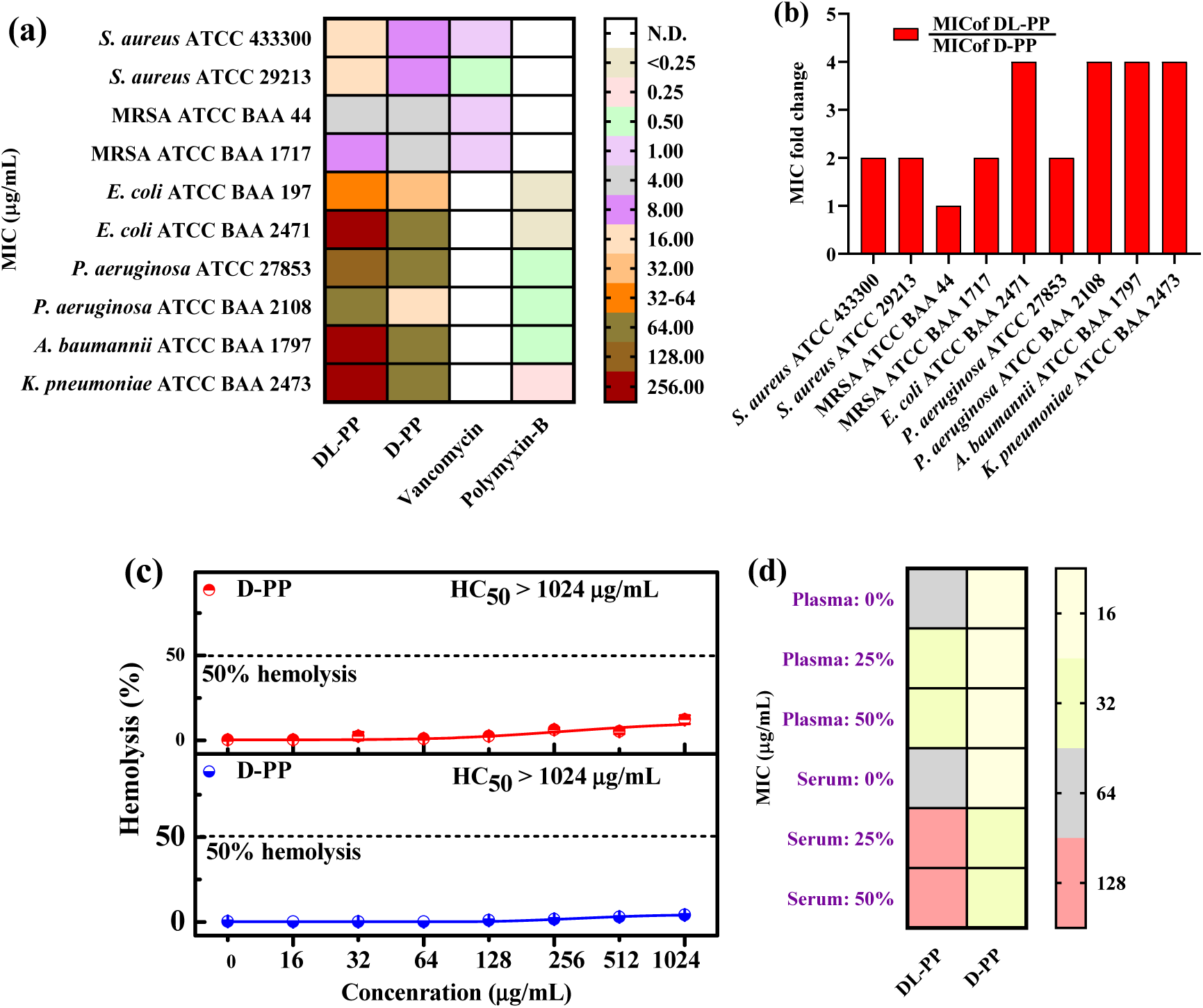
The secondary structure influences antibacterial activity of peptide polymers. (a) Heat-map for antibacterial activity (MIC) of peptide polymers and (b) Comparison of MIC for D-PP vs. DL-PP. Helical peptide polymer outperforms the non-helical; (c) Percentage of hemolysis of mouse erythrocytes by D-PP and DL-PP; and (d) Heat-map representing the antibacterial activity (MIC) of D-PP and DL-PP against *P. aeruginosa* ATCC BAA 2108 upon preincubation with mouse blood plasma and serum.

Given the poor proteolytic stability of typical peptides, both polymers were evaluated and pre-incubated with varying percentages (0, 25, and 50%) of mouse blood plasma and serum, followed by testing against *P. aeruginosa* ATCC BAA 2108. Plasma pre-incubation (25% and 50%) did not change D-PP’s MIC but decreased DL-PP’s MIC by two-fold (Figure 2d). Serum pre-incubation, however, led to a minimal two-fold MIC increase for both polymers. These results suggested that the secondary structure did not influence both polymers’ stability, likely due to the high content of D-amino acids.

Although both polymers effectively targeted Gram-positive bacteria, their limited impact on Gram-negative strains motivated us to evaluate them as adjuvants for antibiotics. Checkerboard assays showed both polymers enhanced antibiotic efficacy by 2-256 folds against multiple strains of bacteria (Figure 3, Tables S1 and S2), highlighting a synergistic or additive effect. Peptide polymers successfully revitalized antibiotics like tetracyclines, ansamycins, fusidanes, macrolides, cephalosporins, and monobactams for Gram-negative bacteria. For instance, MDR *A. baumannii*, typically resistant to fusidic acid, showed a decrease in MIC from 128 µg/mL to 1 µg/mL with only 8 µg/mL of D-PP, achieving a 128-fold potentiation (Figure 3a). D-PP exhibited a two-fold greater potentiation of fusidic acid than DL-PP under the same fractional inhibitory concentration (FIC) of 8 µg/mL.

**Figure 3.**
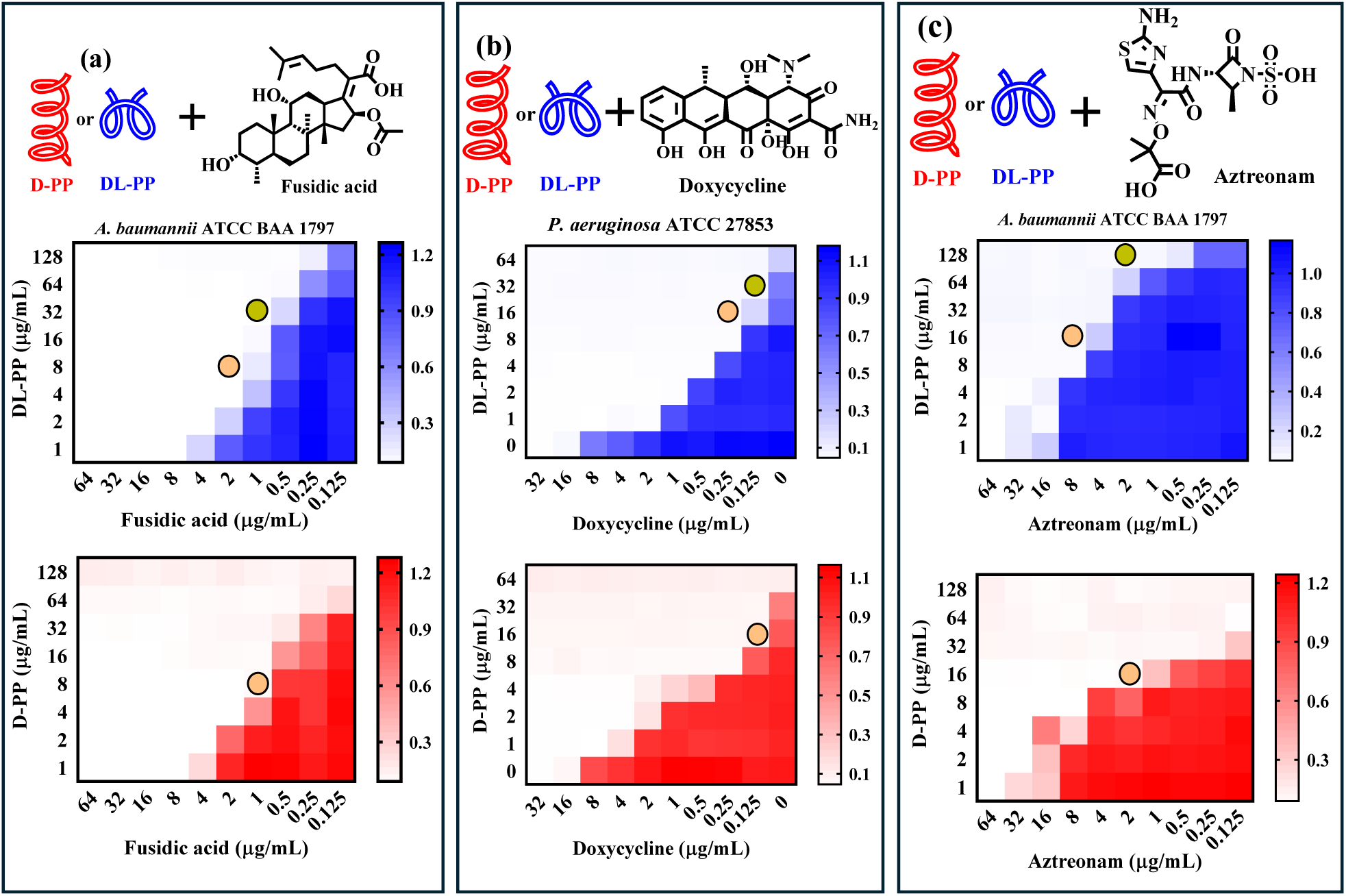
Peptide polymers are universal adjuvants against MDR pathogens. Heatmap represents the checkerboard assays displaying how helical and non-helical peptide polymers at variable concentrations potentiate obsolete antibiotics (e.g. decreased influx, increased efflux and enzymatic degradation). The peptide polymers revitalized (a) fusidic acid (fusidane class), (b) doxycycline (tetracycline class), and (c) aztreonam (monobactam class). The decrease in red or blue colour represents the decrease in microbial cell density.

The polymers also substantially boosted the efficacy of doxycycline and minocycline against *P. aeruginosa*, with D-PP providing a 128-fold enhancement for doxycycline (MIC dropped from 16 to 0.12 µg/mL) at a two-fold lower concentration than DL-PP (Figure 3b). Additionally, D-PP and DL-PP improved the efficacy of aztreonam, commonly affected by β-lactamase with reduced uptake. For instance, D-PP lowered aztreonam’s MIC from 64 µg/mL to 2 µg/mL against MDR *A. baumannii*, achieving 32-fold potentiation, while DL-PP provided 16-fold potentiation at the same concentration (Figure 3c). Three other classes of antibiotics (ansamycins, macrolides, and cephalosporins) were also evaluated for their enhanced efficacy when combined with both peptide polymers, though D-PP consistently exhibited higher potentiation than DL-PP (Figures S8-S12, Tables S1 and S2).

A comparative analysis of the potentiation fold (PF) and fractional inhibitory concentration index (FICI) highlighted the better performance of helical D-PP in potentiating antibiotics.^30^ D-PP achieved 4-fold enhanced potentiation of fusidic acid (PF of 64 to 128) for two *Pseudomonas* strains compared to DL-PP, with synergistic FICIs for both (Figure 4a-4b). For *K. pneumoniae* and *A. baumannii*, D-PP displayed respectively 8-fold and 2-fold higher potentiation than DL-PP. Similarly, D-PP outperformed DL-PP in doxycycline potentiation for *Klebsiella*, *Acinetobacter*, and *Escherichia* species, maintaining synergy across most strains (except for *A. baumannii* where FICI > 0.5 suggested an additive effect) (Figure 4c-4d). Overall, these results confirmed that the helical structure of D-PP enabled more effective potentiation of antibiotics at lower doses compared to DL-PP.

**Figure 4.**
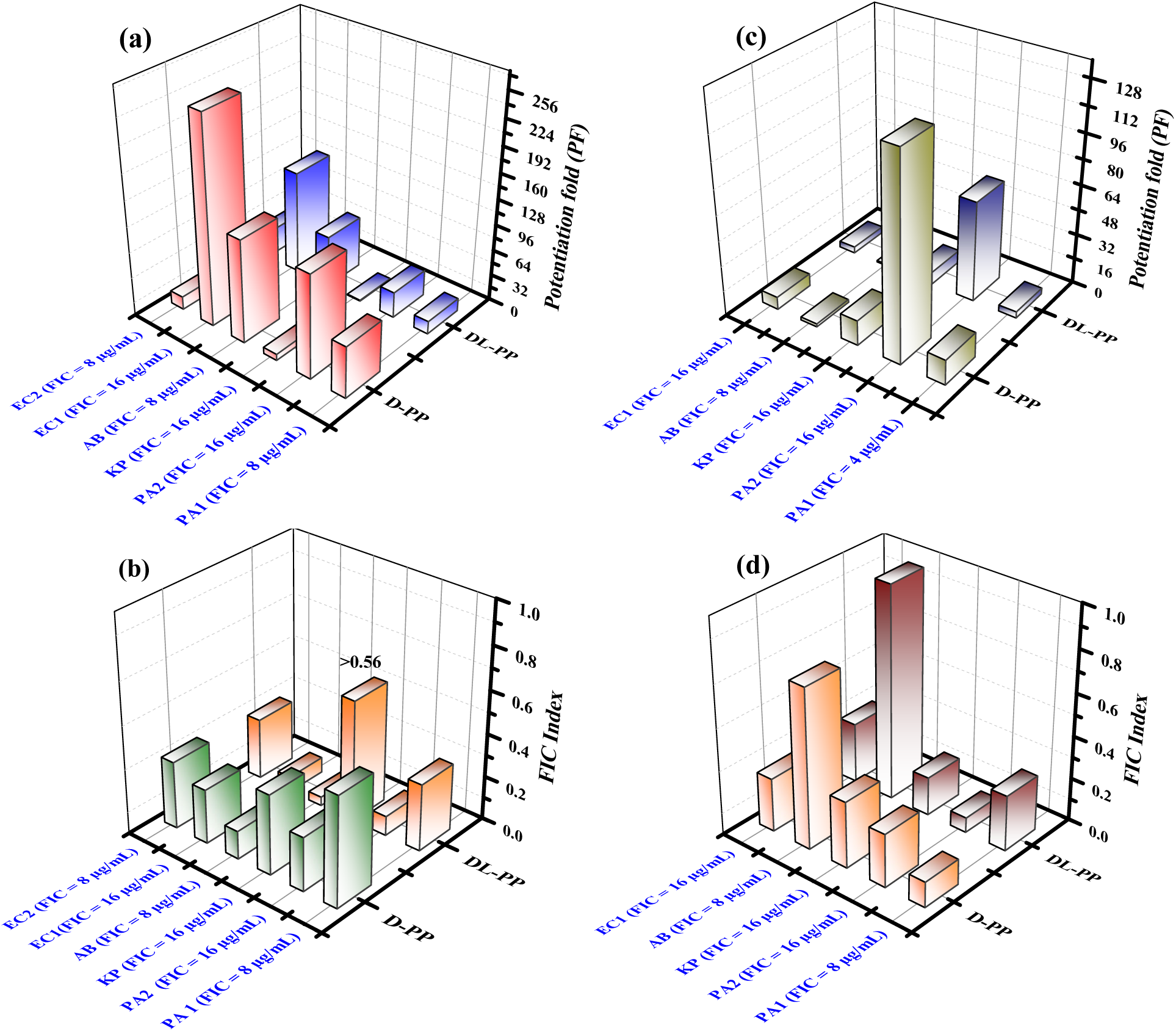
Helical structure favors antibiotic potentiation with lower dosages over non-helical polymer. Fusidic acid and polymer combination: (a) potentiation factor and (b) FICI at a fixed FIC of adjuvant. Doxycycline and polymer combination: (c) potentiation factor and (d) FICI at a fixed FIC of adjuvant. PF stands for potentiation fold indicating the degree to which adjuvants improve the efficacy of antibiotics. FIC refers to the fractional inhibitory concentration of adjuvants required for antibiotic potentiation. FICI represents the fractional inhibitory concentration index of the combinations. FICI = FIC_Adjuvant_ /MIC_Adjuvant_ + FIC_antibiotic_ / MIC_antibiotic._ FICI ≤ 0.5: synergistic, 0.5<FICI<1: additive, 1<FICI<4: no interaction, and FICI>4: antagonism Abbreviations: EC1: *E. coli* ATCC BAA 2741; EC2: *E. coli* ATCC BAA 197; AB: *A. baumannii* ATCC BAA 1797; KP: *K. pneumoniae* ATCC BAA 2473; PA1: *P. aeruginosa* ATCC BAA 2108; and PA2: *P. aeruginosa* ATCC 27853.

In addition, these polymer-antibiotic combinations also demonstrated potent bactericidal effects against planktonic cells, achieving rapid bacterial eradication in MDR Gram-negative *A. baumannii* and *E. coli* through killing kinetics (Figure S17), with no detectable bacterial presence in media or on agar when D-PP was paired with aztreonam or erythromycin. In contrast, treatments with D-PP or antibiotics alone allowed significant survival of these bacteria.

### The helical polymer demonstrated strong interactions with microbial membranes, enhancing both antibacterial activity and antibiotic potentiation

Structurally similar to AMPs, D-PP acted independently against Gram-positive bacteria and effectively targeted Gram-negative bacteria when combined with inactive antibiotics. The standalone antibacterial activity of D-PP was examined through its impact on membrane depolarization using DiSC_3_(5), a dye sensitive to membrane potential changes.^31^ Under normal bacterial membrane conditions, the dye accumulates and quenches its fluorescence; however, membrane disruption causes the dye to leak and the fluorescence to increase. When MRSA ATCC BAA 1717 was treated, D-PP exhibited stronger membrane activity (Figure 5a) at 16 µg/mL (MIC), D-PP increased fluorescence by approximately 20%, whereas DL-PP showed about 5% increase, indicating that D-PP’s helical structure induces more potent membrane depolarization. Similarly, D-PP caused a fluorescent increase when treating *K. pneumoniae* ATCC BAA 2473 than DL-PP (Figure 5b).

**Figure 5.**
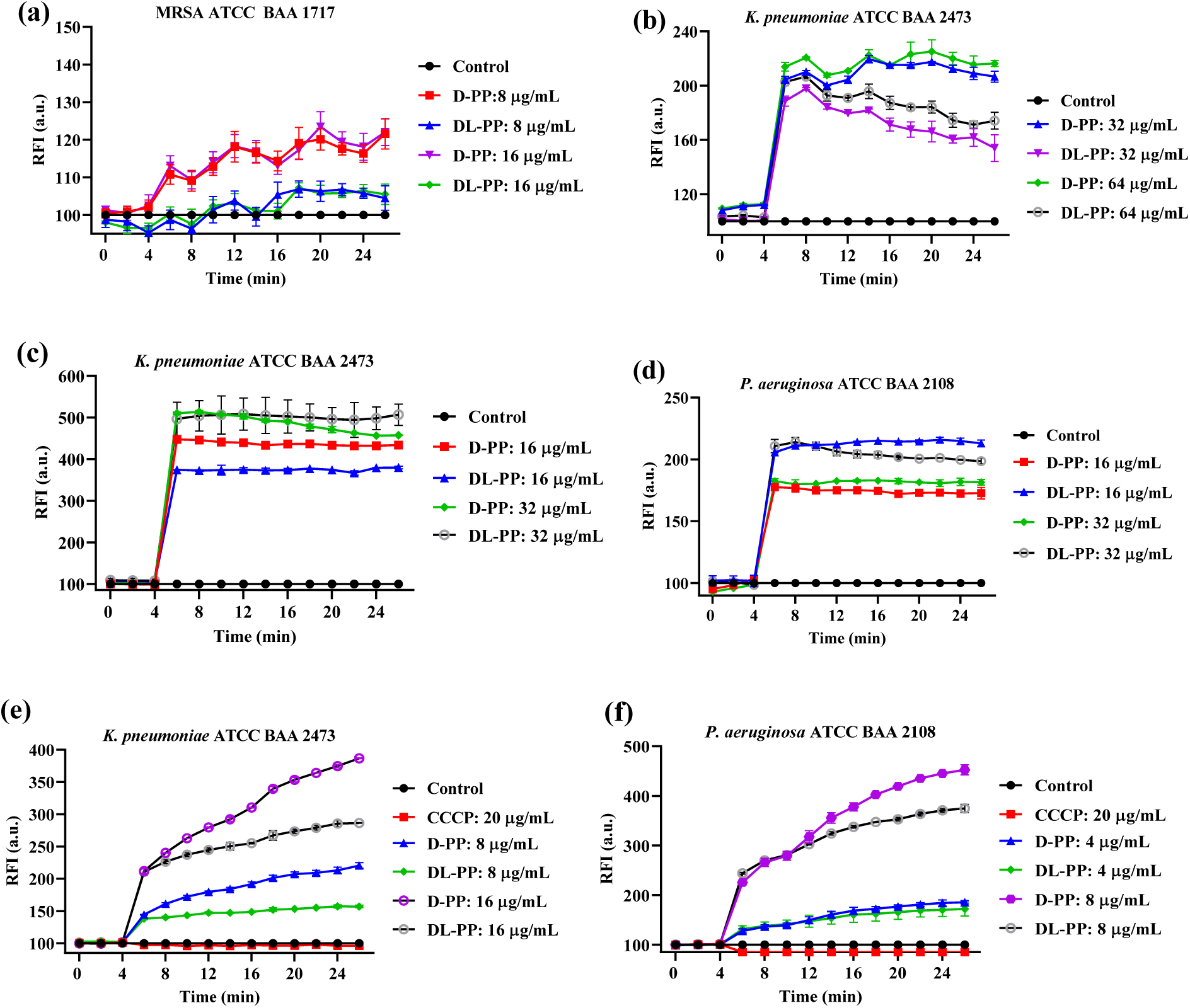
Helical polymer displays enhanced interaction with the microbial membrane over the non-helical polymer, affecting antibacterial activity and antibiotic potentiation efficacy. (a-b): The disruption of bacterial membrane potential by D-PP and DL-PP by monitoring the fluorescence of membrane potential-sensitive dye DiSC_3_(5), (c-d): Monitoring the alteration of bacterial outer membrane potential using fluorescence of dye NPN, (e-f): The analysis of antibiotic uptake (doxycycline) upon treatment with D-PP and DL-PP measuring the fluorescence of the drug itself.

D-PP’s ability to permeabilize bacterial outer membranes, especially in *Klebsiella pneumoniae*, was assessed with *N*-phenyl naphthyl amine (NPN), a hydrophobic dye that fluoresces within the membrane environment.^32^ At 16 µg/mL, D-PP caused a fluorescence increase of ∼65% more than DL-PP in *K. pneumoniae* (Figure 5c), illustrating a stronger outer membrane interaction. This highlighted the outer membrane permeability corresponding to D-PP’s lower FIC when combined with antibiotics, suggesting that D-PP facilitates antibiotic entry by weakening the bacterial outer membrane, particularly for compounds like rifampicin and fusidic acid. However, in *Pseudomonas aeruginosa*, DL-PP caused slightly higher NPN fluorescence (Figure 5d), hinting at different interaction dynamics in this bacterial species, likely due to its non-helical nature. The polymers’ effect on antibiotic uptake within bacterial cells was measured using doxycycline as a model antibiotic. In general, doxycycline fluoresces inside bacterial cells, allowing it to track its intracellular levels.^33^ In both *K. pneumoniae* and *P. aeruginosa*, D-PP facilitated a dose-dependent increase in fluorescence intensity, with levels over 50% higher than DL-PP at the same concentrations (8 or 16 µg/mL) (Figure 5e-5f).

This increased fluorescence suggested that D-PP enhanced doxycycline accumulation by disrupting the bacterial membrane potential (ΔΨ), thus amplifying the ΔpH gradient, a critical component for tetracycline uptake. The experiment was further validated by adding CCCP (carbonyl cyanide m-chlorophenyl hydrazone), known to abolish the PMF across the bacterial membrane (Figure 5e-5f).^34^ CCCP treatment did not increase doxycycline uptake in *P. aeruginosa* and *K. pneumoniae*, as indicated by the very low level of fluorescence, closer to the control.

In addition to membrane permeabilization, D-PP induced intracellular reactive oxygen species (ROS) production in *K. pneumoniae*, which can damage microbial cells by disrupting bacterial homeostasis.^35^ The level of ROS was measured using the cell-permeable dye 2′,7′-dichlorofluorescein diacetate (DCFH-DA), which fluoresces upon reaction with ROS. At 16 µg/mL, D-PP increased ROS levels by 200% more than DL-PP (Figure 6a). Introducing an antioxidant, NAC (N-acetyl cysteine), dramatically reversed the ROS increase, demonstrating that D-PP’s effect on ROS generation significantly contributes to bacterial cell damage. Further, D-PP’s ability to neutralize bacterial surface charge was more pronounced than that of DL-PP (Figure 6b). The surface potential of *K. pneumoniae* shifted from -31 mV to +44 mV after treatment with 1 mg/mL of D-PP, while DL-PP treatment shifted it to +31 mV. The surface potential moved to +20 mV and +8 mV after being exposed to 64 µg/mL of helical (D-PP) and non-helical (DL-PP) polymers respectively. The more substantial shifts with D-PP suggest that the helical structure more effectively disrupts the bacterial cell envelope, which could destabilize bacterial growth and metabolism by altering surface potential.

**Figure 6.**
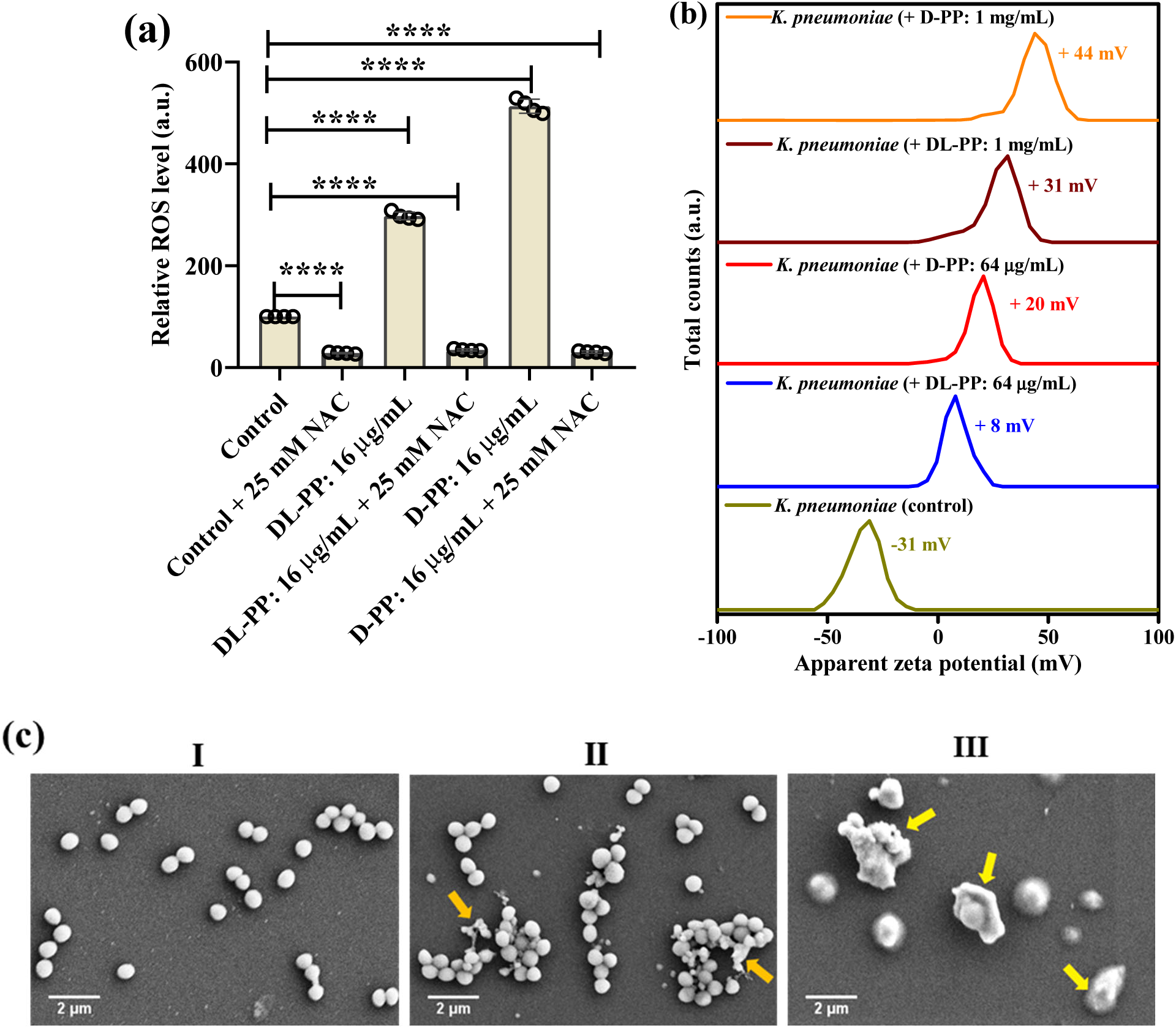
The helical polymer outperformed its non-helical in interacting with bacteria. (a) ROS accumulation in *K. pneumoniae* ATCC BAA 2473 upon exposure to peptide polymers whereas the addition of an antioxidant, NAC, diminished the ROS level; (b) Alteration of Zeta potential of bacterial cell (*K. pneumoniae* ATCC BAA 2473) suspension with the treatment of D-PP and DL-PP; (c) SEM images of MRSA ATCC BAA 1717: (I) Untreated, (II) Treated with DL-PP (16 µg/mL), and (III) Treated with D-PP (16 µg/mL). Statistical analysis was conducted using GraphPad Prism 7 software, employing ordinary one-way ANOVA with Dunnett’s multiple comparisons test (***: P <0.0001).

The structural damage induced by D-PP was visualized through scanning electron microscopy (SEM) (Figure 6c). At 16 µg/mL, D-PP-treated MRSA cells showed marked morphological alterations, with more irregular shapes and a notable increase in cellular debris compared to DL-PP treatment. The untreated MRSA cells maintained a healthy circular shape, whereas the D-PP-treated cells exhibited distorted shapes and membrane debris clusters, particularly evident with D-PP treatment, implying greater membrane damage from the helical polymer.

Combining all these results, it is evident that a helical conformation promotes more effective interactions with bacterial cells, thereby enhancing antimicrobial efficacy compared to a random, non-helical structure. Notably, the DL-polypeptide also exhibits valuable potential, demonstrating potency against some pathogens. While prior studies have scarcely investigated the pronounced differences in cell penetration,^6,17-19^ the findings in this work represent a significant advancement in the design of both D- and DL-peptide polymers. Given the overall superiority of the helical D-PP, the following sections further examine its impact on genomic alterations, its efficacy against dormant sub-populations and biofilms of pathogenic bacteria, and its potential to resist microbial adaptation.

### The helical polymer adjuvant affects bacterial membrane stability and metabolism through transcriptional regulation

Bacteria commonly initiate metabolic adjustments to maintain cellular stability in response to external stressors. These responses are essential for enhancing antibiotic efficacy. To investigate the transcriptional shifts, transcriptomic analysis was conducted in *E. coli* and *K. pneumoniae* treated with D-PP at concentrations of 8 and 16 µg/mL for 6 hours. This analysis, based on BioCyc Genome Database Collection,^36^ revealed upregulation and downregulation patterns among differentially expressed genes (DEGs) related to outer and inner membrane structures, as well as cellular exporters and importers—key factors for potentiating antibiotic effects (Figure 7 and Tables S3-S6). For instance, in *E. coli*, D-PP treatment led to upregulation of the inner membrane protein gene *YaiY*, signaling disruptions to bacterial inner membrane function. Additionally, there was an overexpression of *OmpX* gene for the outer membrane protein, known for its roles in maintaining membrane stability, integrity, permeability, and pathogenicity. Moreover, D-PP promoted the upregulation of the ABC transporter ATP-binding subunit YbhF, which can facilitate antibiotic resistance by enabling the efflux of antibiotics like tetracyclines through ATP-driven processes (Figure 7a). In *K. pneumoniae*, the polymer adjuvant-induced overexpression of the gene for envelope stress response proteins *PspG*, a crucial factor for managing membrane integrity and permeability stress. Additionally, downregulation was observed in *K. pneumoniae* for genes encoding ABC transporters involved in nutrient uptake, such as ribose, glycerol-3-phosphate, and dipeptides (*RbsA, UgpC,* and *DppC*) (Figure 7b). These transporters contribute directly or indirectly to virulence and antibiotic resistance. Collectively, these results supported previous mechanistic findings that D-PP disrupts bacterial inner and outer membranes and interferes with efflux systems, thereby enhancing the efficacy of antibiotics that are typically limited by membrane permeability and efflux barriers. The adjuvant also aided in reducing resistance factors associated with bacterial transport systems.

**Figure 7.**
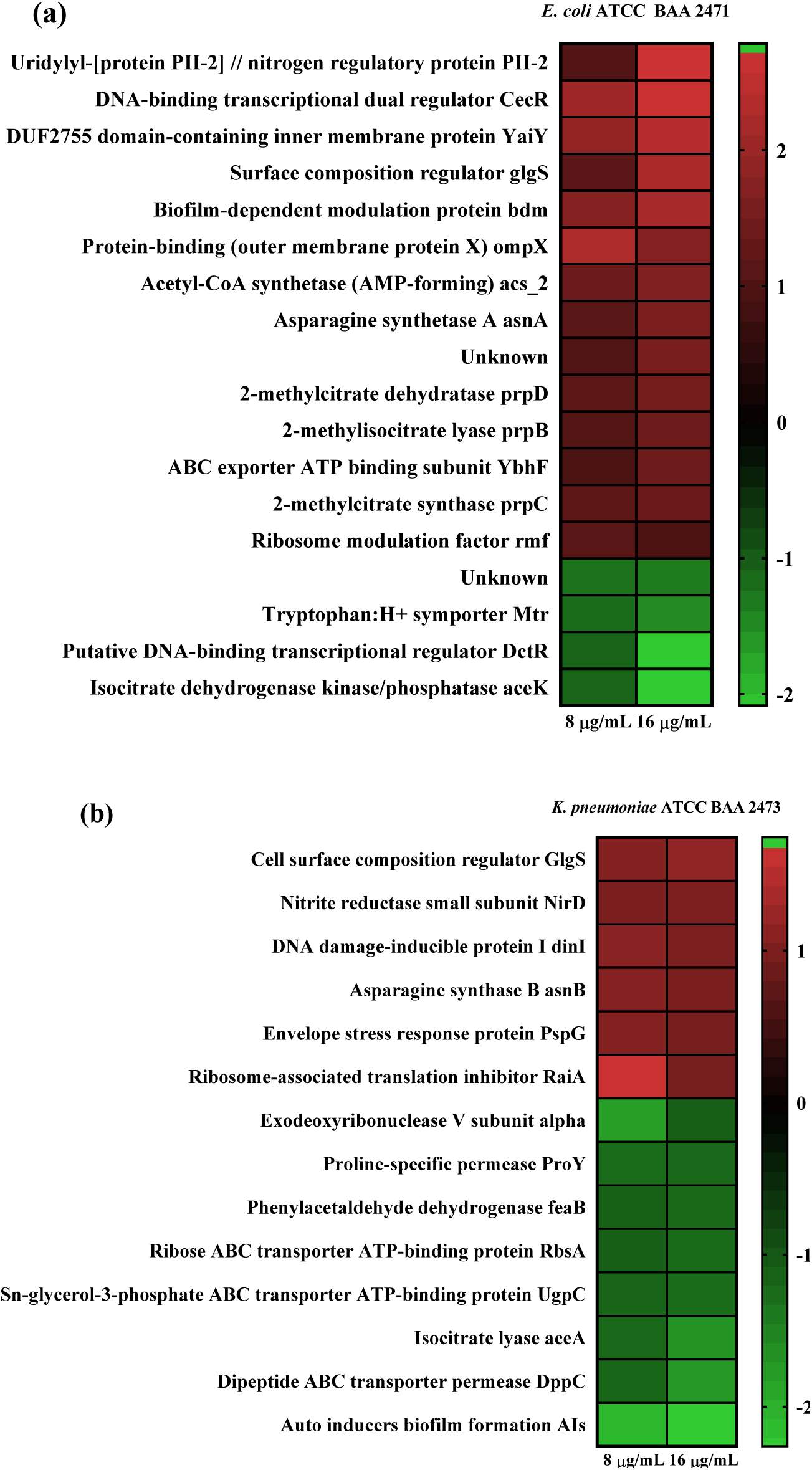
Metabolic alterations in bacterial cells by the helical peptide polymer (D-PP). Heatmap represents the transcriptomic analysis of (a) *E. coli* and (b) *K. pneumoniae* upon incubation of D-PP at 8 and 16 µg/mL for 6 h. Differentially expressed genes (DEGs) were identified between the treatment of D-PP (8-16 µg/mL) vs. untreated control with a Log_2_(Fold change) ≥ 1 and Log_2_(Fold change) ≤ -1 and P value < 0.05. The DEGs were selected in the heatmap from the original list of individual treatments vs. control (Tables S3-S6), if they are co-upregulated or co-downregulated in both treatment conditions. All data are represented as the means of three biological replicates.

**The helical polymer adjuvant-antibiotic combinations showed strong bactericidal effects against dormant bacterial cells,** providing alternative therapeutics to address recurrent and chronic infections associated with these sub-populations, beyond common planktonic cells. Dormant bacteria, typically found in the stationary phase of growth, shut down essential cellular processes like cell division and protein synthesis, making them resistant to conventional antibiotics targeting these processes.^37-39^ The efficacy of D-PP and antibiotic combinations over time (0, 4, 8, and 24 hours) was assessed against dormant populations of various MDR Gram-negative pathogens (Figure 8a-8b and Figure S16). For instance, D-PP (16 µg/mL) with erythromycin (16 µg/mL) gradually reduced stationary *E. coli* cells over time, achieving ∼2-log (99%) reduction in 24 hours. Increasing erythromycin to 32 µg/mL with the same D-PP concentration improved reduction to 3.3-log (99.9%) by 24 hours (Figure 8a). In the case of *K. pneumoniae*, the combination of D-PP (32 µg/mL) and doxycycline (2–4 µg/mL) achieved complete eradication (∼6.6 log reduction) of dormant cells within 24 hours (Figure 8b). It was surprising to observe that D-PP alone at the same concentration was similarly effective against dormant cells of *K. pneumoniae*, a stark contrast to its limited efficacy against planktonic cells. Doxycycline alone showed no effect, and polymyxin-B alone resulted in only a 1-log reduction. Likewise, the D-PP (16 µg/mL) combined with aztreonam (2–4 µg/mL) quickly eliminated stationary cells of *A. baumannii* within 2 hours, unlike standalone treatments with aztreonam or polymyxin-B (Figure S16). These findings underscored the potent bactericidal nature of D-PP-antibiotic combinations against metabolically inactive, dormant bacterial sub-populations.

**Figure 8.**
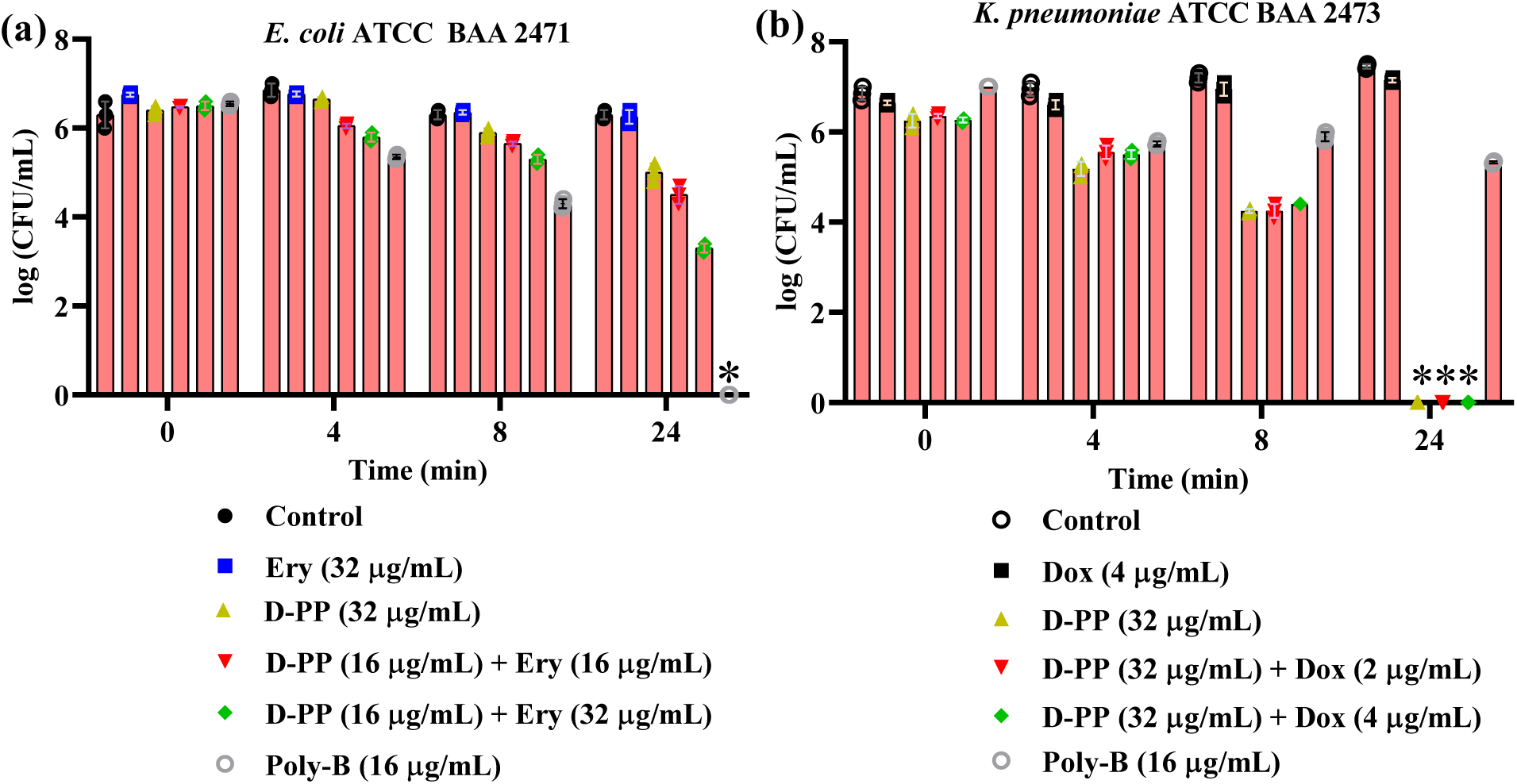
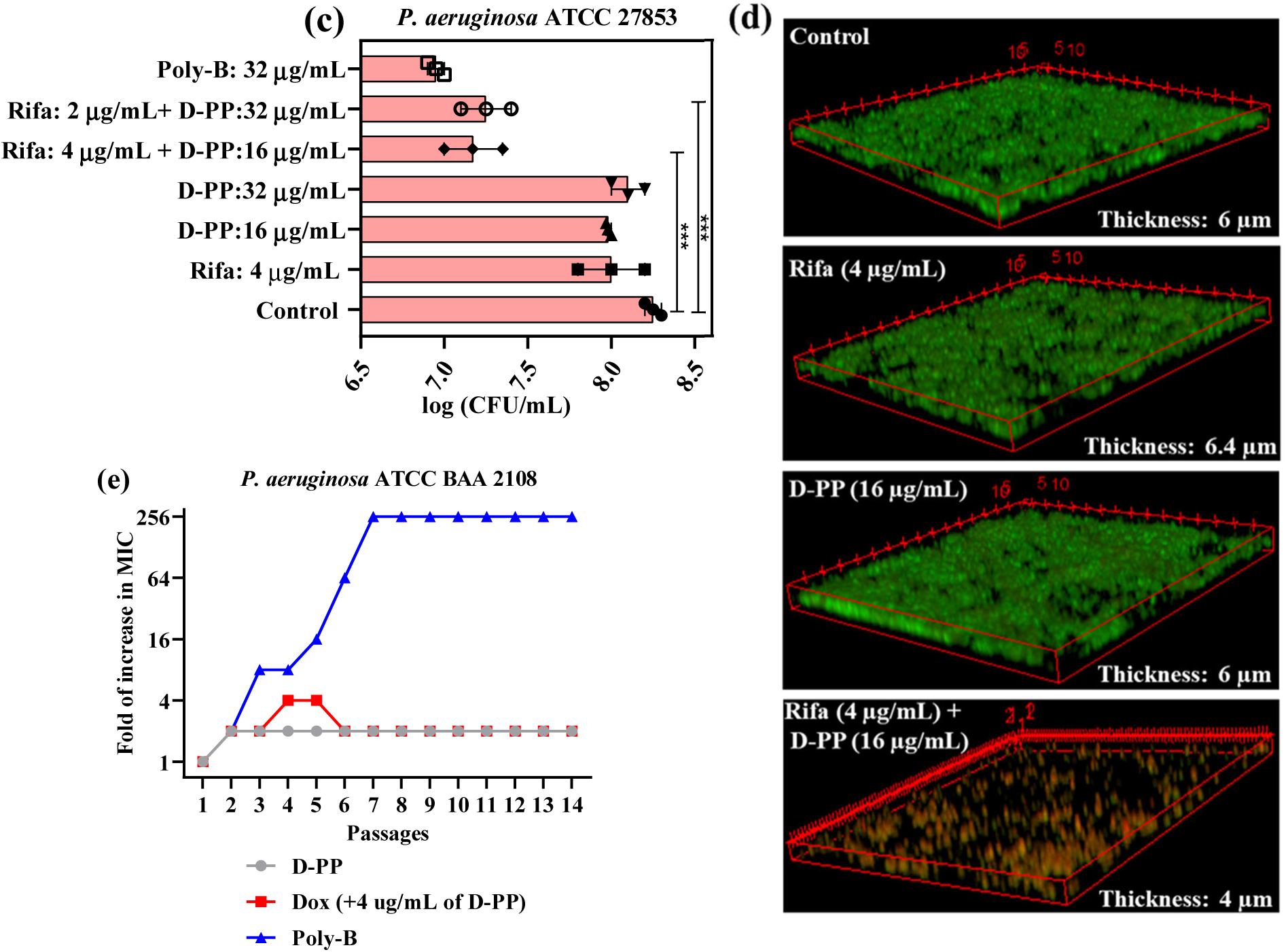
Efficacy of the helical peptide polymer (D-PP) and antibiotic combinations against dormant cells and biofilms with low resistance propensity. Bactericidal kinetics against dormant sub-populations with (a) D-PP and erythromycin (Ery) combination; and (b) D-PP and doxycycline (Dox) combination. *Pseudomonas* biofilm disruption by the treatment of D-PP and rifampicin (Rifa) combination: (c) Cell viability of biofilm-embedded bacteria and (d) Confocal fluorescence microscopy to visualize biofilm disruption. (e) Bacterial (*P. aeruginosa* ATCC BAA 2018) resistance development against D-PP standalone, polymyxin-B (Poly-B) and Dox-D-PP combinations. Statistical analysis was conducted using GraphPad Prism 7 software, employing ordinary one-way ANOVA with Dunnett’s multiple comparisons test (**P = 0.0016 and ***P = 0.0003).

### The helical polymer adjuvant-antibiotic combinations showed strong efficacy against bacterial biofilms and demonstrated a low propensity for resistance development

Biofilms consist of extracellular polymeric substances (EPS) that form a barrier to antibiotics. This EPS, composed of negatively charged components like polysaccharides, proteins, and nucleic acids, shelters diverse bacterial populations in a protected environment, making conventional antibiotics largely ineffective against biofilm-associated infections.^39-41^ As a model study, D-PP and rifampicin combinations were tested their effects on mature *Pseudomonas* biofilms (Figure 8c). The combination of D-PP (16 µg/mL) with rifampicin (4 µg/mL) achieved a 94% reduction (∼1.2 log) in biofilm-embedded bacteria. Doubling the D-PP concentration and reducing rifampicin to 2 µg/mL yielded similar results, with a ∼90% reduction (∼1 log). Polymyxin-B, often used as a last-resort antibiotic for Gram-negative infections, produced comparable results (∼90% reduction), but D-PP or rifampicin alone had a minimal effect on biofilm-associated bacteria. Confocal fluorescence microscopy provided further qualitative analysis (Figure 8d). Dual staining with SYTO-9 (green for live/compromised cells) and propidium iodide (red for dead cells) confirmed that the D-PP-rifampicin combination reduced biofilm thickness from 6 µm in the untreated control to 4 µm, with substantial bacterial cells being dead. In contrast, D-PP or rifampicin alone, as well as untreated samples, retained thick biofilms (6–6.4 µm) with most cells being live. The amphiphilic structure of D-PP, featuring pendant ammonium residues, likely facilitates interactions with EPS’s negatively charged components, enhancing antibiotic penetration and enabling the effective killing of biofilm-embedded bacteria.

We further tested whether the helical D-PP could prevent resistance to doxycycline over 14 passages of bacterial exposure. Alongside D-PP-doxycycline, we included standalone D-PP and polymyxin-B as controls (Figure 8e). Initially, the MIC of doxycycline was 2 µg/mL (with D-PP at 4 µg/mL) against MDR *P. aeruginosa*, while the standalone MICs for D-PP and polymyxin-B were 16 µg/mL and 0.5 µg/mL respectively. Over four bacterial passages, doxycycline’s MIC in the D-PP combination increased by only 4-fold, stabilizing at a 2-fold increase at passage 14, showing minimal resistance development. Similarly, D-PP alone showed only a 2-fold MIC increase through the entire passages. In stark contrast, polymyxin-B displayed an 8-fold MIC increase after only 3 passages, reaching a 256-fold increase by 7 passages, which persisted throughout the experiment, indicating rapid and significant resistance development. Together the D-PP-antibiotic combination demonstrated superior biofilm eradication and minimized resistance development compared to polymyxin-B, underscoring its potential as a promising candidate for durable antimicrobial therapy.

**In conclusion,** this study provides critical insights about the effects of secondary structures on mechanisms of action and antimicrobial efficacy in synthetic AMP mimics. The dextrorotatory peptide polymer (D-PP) adopts a well-defined α-helix under biomimetic conditions. In contrast, the mixed dextrorotatory-levorotatory polymer (DL-PP) shows a random conformation without a defined secondary structure. This structural distinction is crucial: The helical structure in D-PP increases antibacterial potency against Gram-positive bacteria, making it more effective than the non-helical DL-PP while maintaining similar hemolytic activity and protease stability. Beyond its standalone antimicrobial effects, D-PP restored the efficacy of various classes of antibiotics against Gram-negative bacteria, with 2 to 256 potentiation folds. Notably, D-PP achieved better antibiotic potentiation at lower doses than DL-PP, underscoring the role of helical structures in optimizing both antibacterial and adjuvant capabilities. Mechanistic studies revealed D-PP’s stronger interaction with bacterial membranes compared to DL-PP, establishing D-PP’s amphiphilic helical structure as the preferred therapeutic candidate. While D-PP stands out for its potent antibacterial and adjuvant properties, DL-PP could be useful as an effective, lower-cost alternative against some pathogens, particularly in resource-limited settings where cost is a critical factor.

## EXPERIMENTAL METHODS

The major experimental methods are depicted in this section and the remaining protocols for biological assays are described in the supplementary information.

### Synthesis of D/DL-allylglycine NCA.^26^

D or DL-allylglycine (1.0 g, 8.7 mmol, 1 eq.), 50 mL of tetrahydrofuran (THF), and 6.2 mL of propylene oxide (87.5 mmol, 10 eq.) were sequentially added to a pressure vessel. Further, 1.3 g of triphosgene (4.4 mmol, 0.5 eq.) was carefully added and the mixture was stirred vigorously with a magnetic stir bar for 20 h at room temperature. After cooling on ice for an hour, 15 mL of water was added to the mixture to quench any unreacted triphosgene for safety. After 5-10 min of stirring, THF was removed completely from the reaction mixture using a Rotary evaporator. Finally, ethyl acetate was added to the mixture and the organic phase was washed thrice with brine solution. Next, the organic phase was concentrated, and the product was purified through crystallization using a THF and hexane mixture at <10 °C, yielding a white crystalline product (yield: 80-85%).

DL-allylglycine NCA: FT-IR (cm^-1^): 3347 (amide N-H Str.), 1828 (anhydride C=O asym. Str.), 1756 (anhydride C=O sym. Str.), 1630 (amide I C=O Str.), 1520 (amide II C=O str.); ^1^H-NMR (400 MHz, DMSO-d_6_): δ/ppm. 9.1 (bs, -N*H*CO, 1H), 5.6-5.8 (m, CH_2_C*H*CH_2_-, 1H), 5.1-5.2 (m, C*H_2_*C*H*CH_2_-, 2H), 4.5 (t, -C*H*_(Allylglycine)_, 1H), 2.4-2.5 (m, -CHC*H_2_*CH-, 2H). ^13^C-NMR (100 MHz, DMSO-d_6_): 171, 151.9, 131.4, 120, 56.9 and 34.9. HR-MS (m/Z): 141.0420 [M^+^] (observed), 141.0426 [M^+^] (calculated).

D-allylglycine NCA: FT-IR (cm^-1^): 3288 (amide N-H Str.), 1826 (anhydride C=O asym. Str.), 1770 (anhydride C=O sym. Str.), 1643 (amide I C=O Str.), 1515 (amide II C=O str.); ^1^H-NMR (400 MHz, DMSO-d_6_): δ/ppm. 9.1 (bs, -N*H*CO, 1H), 5.6-5.8 (m, CH_2_C*H*CH_2_-, 1H), 5.1-5.2 (m, C*H_2_*C*H*CH_2_-, 2H), 4.5 (t, -C*H*_(Allylglycine)_, 1H), 2.4-2.5 (m, -CHC*H_2_*CH-, 2H). ^13^C-NMR (100 MHz, DMSO-d_6_): 171, 151.9, 131.4, 120, 56.9 and 34.9. HR-MS (m/Z): 141.0436 [M^+^] (observed), 141.0426 [M^+^] (calculated).

### Synthesis of poly(D/DL-allylglycine) via ROP (M/I= 30)

DL-allylglycine NCA monomer (250.0 mg, 1.8 mmol, 30 eq.) was dissolved with 2 mL dry dimethyl formamide (DMF) in a 10 mL Schlenk flask. After three freeze-pump-thaw cycles of the monomer solution, 12.9 µL of benzylamine (0.06 mmol, 1 eq.) was injected and stirred vigorously at room temperature for 24 h in a nitrogen atmosphere. Finally, the reaction mixture was precipitated with diethyl ether to obtain the white lumpy polymer which was characterized through proton-NMR-based end group analysis and FT-IR. For synthesizing poly(D-allylglycine), the same protocol was followed, except the D-allylglycine monomer was dissolved in a 1:1 mixture of dry THF and DMF. The polymers were characterized by FT-IR and NMR (Figures S1 and S2).

Poly(DL-allylglycine) FT-IR (cm^-1^): 3285 (amide N-H Str.), 3078 (aromatic C-H Str.), 1626 (amide I C=O Str.), 1510 (amide II C=O Str.); Poly(D-allylglycine) FT-IR (cm^-1^): 3283 (amide N-H Str.), 3076 (aromatic C-H Str.), 1626 (amide I C=O Str.), and 1520 (amide II C=O Str.).

### Synthesis of D/DL-peptide polymer via thiol-ene click reaction

Poly(D/DL-allylglycine) (50.0 mg, 1 eq.), cysteamine hydrochloride (585.6 mg, 10 eq.), and 2,2-dimethoxy-2-phenyl acetophenone (DMPA) (26.4 mg, 0.2 eq.) were mixed and dissolved in a 10 mL Schlenk flask using 1:1 mixture of DMF and water. After three freeze-pump-thaw cycles, the reaction mixture was stirred vigorously at room temperature under 365 nm UV light. After the first 24 h, 13.2 mg of DMPA dissolved in 0.5 mL DMF was injected into the reaction mixture and continued stirring for the next 48 hours. The final polymer was isolated by vigorous dialysis across the water using 0.5-1 KDa dialysis membrane for 3 days followed by lyophilization. The peptide polymers (D-PP and DL-PP) appeared as white fluffy solids and were characterized through FT-IR and NMR (Figures S1-S8).

DL-PP FT-IR (cm^-1^): 3272 (amide N-H Str.), 3046 (aromatic C-H Str.), 2920 (amine N-H Str.) 1696, 1627 (amide I C=O Str.), 1536 (amide II C=O Str.); D-PP FT-IR (cm^-1^): 3272 (amide N-H Str.), 3037 (aromatic C-H Str.), 2918 (amine N-H Str.) 1651, 1624 (amide I C=O Str.), 1516 (amide II C=O Str.).

### Secondary structure characterization

Circular dichroism (CD) spectroscopy was employed to assess secondary structures of polymers. The polymers were dissolved in different solvents at 0.25 mg/mL and incubated the solution at 37 °C for different time depending upon the solvents. CD spectra were recorded with 0.2 mL of polymer solutions using a quartz cuvette (path length: 1 mm) at the wavelength range of 190-250 nm. Poly(D/DL-allylglycine) was dissolved in 100% HFIP. D-PP and DL-PP were dissolved in 1×PBS and incubated for 3 days. The peptide polymers (D/DL-PP) were also dissolved in a 1:1 mixture of water and HFIP (a biomimicking environment).

### Bacterial culture conditions

All the drug-sensitive and multidrug-resistant bacterial strains (Figure 2a) were purchased from ATCC. In general, the bacteria from their primary glycerol stocks were streaked on the solid tryptic soy agar (TSA). After 24 h incubation at 37 °C, a single bacterial colony from the TSA plate was inoculated into a tryptic soy broth (TSB) at 37 °C for 6 h under constant shaking at 190 rpm. This resulted in the production of mid-log or exponential-phase bacteria. For certain bacterial stains, specific antibiotics were used in solid agar and liquid broth. For example, ceftazidime (10 µg/mL) was for *E. coli* ATCC BAA 197 and imipenem (25 µg/mL) was for *E. coli* ATCC BAA 2471 and *K. pneumoniae* ATCC BAA 2471.

### Antibacterial assay

The peptide polymers were 2-fold serially diluted using media [1:1 cation adjusted Muller Hinton Broth (CMHB) and normal saline (NS)] from the highest-tested concentration in a 96-well plate. Further, 150 µL of the exponential phase bacterial suspension (∼10^5^ CFU/mL) in the same media was added to all the wells containing 50 µL solution of the polymer at different concentrations. The whole plate was incubated at 37 °C under shaking for 16-18 h and optical density (O.D.) at 600 nm was recorded using a microplate reader. Each concentration was in triplicates and the lowest concentration of the test drug showcasing OD <0.1 was considered as the minimum inhibitory concentration (MIC).

### Antibacterial activity in plasma and serum

250 µL of a polymer (D/DL-PP) solution in saline (1024 µg/mL) was incubated with 250 µL of 100% and 50% mouse blood plasma and serum for 3 h at 37 ° C under shaking. After the incubation, the polymer solution was used to determine the MIC against MDR *P. aeruginosa* ATCC BAA 2108 by following the above-mentioned protocol of antibacterial assay.

### Antibacterial activity of combinations

It was evaluated through checkerboard assay. The polymer was serially 2-fold diluted from the highest concentration using 1:1 CMHB and NS in a 96-well plate along the row. Next, 25 µL of the 2-fold serially diluted test antibiotic solution was added to the wells containing 25 µL of polymer solution of variable concentration. Further, 150 µL of mid-log phase bacterial suspension (∼10^5^ CFU/mL) was added to the 50 µL mixture of polymer and antibiotic. After 16-18 h incubation at 37 °C, OD at 600 nm was recorded using a microplate reader. The FIC of the antibiotic was determined at the various sub-MIC concentrations of the polymer, showcasing OD <0.1.

### Bacterial membrane depolarization assay

A mid-log phase bacterial suspension (∼10^8^ CFU/mL) was centrifuged at 3500 rpm for 5 min and the bacterial palate was resuspended in a 1:1:1 ratio of 5 mm glucose, 5 mm HEPES, and 100 mm KCl with 2 µM dye DiSC_3_(5) (5,3,3′-dipropylthiadicarbocyanine iodide). Additionally, the suspension was supplemented with 0.2 mM ethylenediaminetetraacetic acid (EDTA) for Gram-negative bacteria. The bacterial suspension with dye was incubated at dark at least for an hour. Subsequently, 180 µL of the suspension was transferred to a black 96-well plate and the fluorescence of DiSC_3_(5) was monitored at an excitation wavelength of 622 nm and an emission wavelength of 670 nm for 2 min intervals. After 4 min, 20 µL of the polymer’s aqueous solution was added to the well and fluorescence monitoring continued till 24 min. Each treatment and the control had triplicate values in the study.

### Outer membrane permeabilization assay

A mid-log phase bacterial suspension (∼10^8^ CFU/mL) was centrifuged at 3500 rpm for 5 min and the bacterial palate was resuspended in a 1:1 ratio of 5 mm glucose and 5 mm HEPES with 10 µM dye NPN. Subsequently, 180 µL of the dye-containing suspension was transferred to a black 96-well plate and the fluorescence of NPN was monitored at an excitation wavelength of 350 nm and an emission wavelength of 420 nm for 2 min intervals. After 4 min, 20 µL of the polymer’s aqueous solution was added to the well and fluorescence monitoring continued till 24 min. Each treatment and the control had triplicate values in the study.

### Antibiotic uptake assay

A mid-log phase bacterial suspension (approximately 10^8^ CFU/mL) was centrifuged at 3500 rpm for 5 minutes, and the resulting bacterial pellet was resuspended in 10 mM HEPES with 128 µg/mL of doxycycline. Then, 180 µL of this doxycycline-enriched suspension was added to a black 96-well plate, where the fluorescence of doxycycline was measured at an excitation wavelength of 405 nm and an emission wavelength of 535 nm in 2-minute intervals. After 4 minutes, 20 µL of the polymer’s aqueous solution was introduced, and fluorescence readings continued for an additional 24 minutes. 20 µL water and CCCP solution were added instead of the polymer as an untreated and negative control. Each treatment and the control had triplicate values in the study.

### ROS generation

Briefly, 1 mL of bacterial suspension (mid-log phase, ∼10^8^ CFU/mL) in normal saline was incubated with 10 µL of polymer solution (16 µg/mL), either with or without antioxidant, NAC (25 mM) for 2 h at 37 °C under continuous shaking at 180 rpm. Next, 5 µL dye DCFH-DA (5 µM) was added to the treated and untreated bacterial suspension and incubated for another 30 min at dark. The bacterial suspension was further centrifuged at 3500 rpm for 5 min to remove the polymer and the external dye. Next, after resuspension of the bacterial palate with saline, the fluorescence intensity of the dye was recorded at 488 nm excitation and 525 nm emission wavelengths. The experiment was performed in triplicate.

### Transcriptomic analysis.^42^

The mid-log phase bacterial suspension (1 mL of 10^8^ CFU/mL) in 1:1 cationic adjusted Muller Hinton Broth and normal saline was treated with 8 and 16 µg/mL of D-PP for 6 h at 37 °C under shaking (190 rpm). The control bacterial sample was not subject to polymer treatment. The treated and untreated bacterial suspensions were centrifuged for 5 min at 3500 rpm and washed with normal saline. Finally, the bacterial palates were resuspended with 1 mL of DNA/RNA shield lysis buffer. The bacterial lysate was next transferred to a silica bead containing a 2 mL tube and vortexed for an hour. Next, the bacterial samples were subjected to RNA extraction according to the manufacturer’s protocol (Zymo Quick-RNA Fungal/Bacterial Kits, Zymo Research, Irvine, CA). After a purity check of RNA extract quality using nanodrop and Agilent 5200 Fragment Analyzer (Agilent), RNA libraries were prepared using NEBNext® rRNA Depletion Kit (Bacteria), NEBNext Ultra II Directional Library Prep Kit and NEBNext® Multiplex Oligos for Illumina® (NEB, Lynn, MA). Each library was subject to sequencing (Illumina NovoSeq X, S4 flow cell, 150 bp, pair-ended, Psomagen Rockville, MD). Sequences were aligned to *Escherichia coli* (ATCC BAA-2471) and *Klebsiella pneumoniae* (ATCC BAA-2473) using STAR v.2.7.10b. Only those reads that mapped uniquely to the bacteria genome using the annotation file (Escherichia_coli_ATCC_BAA_2471.gbk, Klebsiella_pneumoniae_ATCC_BAA_2473.gbk) were used for gene expression analysis. The expression profile of the significantly expressed genes (cpm > 1) underwent principal component analysis (PCA). For differentially expressed gene (DEG) analysis, the counts data were normalized using the median-of-ratios method and differential analysis performed (DESeq2 pipeline). The average reading depth for the samples was around 20 million reads per sample. RNA-Seq data from this study are available in the GEO database.

### Bacterial biofilm disruption assay

Biofilm disruption experiments were carried out using 18 mm glass coverslips. Initially, a 6-hour culture of *P. aeruginosa* in the mid-log phase was diluted to ∼10^5^ CFU/mL in nutrient broth, which was supplemented with 1% glucose and 1% NaCl. Following this, 2 mL of the diluted bacterial suspension was added to wells containing sterilized coverslips, and the plate was incubated statically at 30 °C for 72 hours. After the media was removed, the biofilm-coated coverslips were gently washed with 1×PBS (pH 7.4) to eliminate any remaining planktonic cells and then transferred to a new 6-well plate. Subsequently, 2 mL of solutions containing D-PP (32 µg/mL), rifampicin (4 µg/mL), and their combinations (rifampicin: 2 µg/mL and D-PP: 32 µg/mL; rifampicin: 4 µg/mL and D-PP: 16 µg/mL) were added to the wells with biofilm-coated coverslips and incubated for 24 hours. As a control, 2 mL of fresh media without any compounds was added to a separate well. Polymyxin-B at 32 µg/mL served as the antibiotic control for this study. After 24 hours, the coverslips were washed with 1×PBS to remove planktonic cells. The washed coverslips containing biofilm were then placed into a fresh well plate, and 2 mL of a trypsin-EDTA solution diluted in saline (1:4) was added, followed by incubation for 10-15 minutes with shaking. A 20 μL sample of the resulting bacterial suspension was then serially diluted tenfold and each dilution was spot-plated on nutrient agar plates, which were incubated for 24 hours. After incubation, viable bacterial colonies were counted, and results were reported as Log CFU per mL.

In a separate experiment, both treated and untreated coverslips from the biofilm disruption assays were washed with 1×PBS and placed on glass slides. Biofilm staining was conducted using a mixture of 5 µL Syto-9 (60 µM) and propidium iodide (PI) (15 µM), and images were captured using a Zeiss LSM 410 Confocal Laser Scanning Microscope. Image processing was performed using Image J software.

### Statistical analysis

Data presented as mean ± standard deviation. Comparisons among different groups in animal studies were conducted using ordinary one-way ANOVA with Dunnett’s multiple comparisons test. A p-value < 0.05 was considered significant. Statistical analysis was performed using GraphPad Prism 7 software.

## Acknowledgements

We acknowledge the National Institutes of Health (R01AI149810) for partial support of our antimicrobial research. We also thank the Functional Genomics Core of the University of South Carolina (supported by P30GM154632) for their help in transcriptomic study. We also thank Dr. Hua Lu from Peking University for the fruitful discussion about polymer synthesis and Dr. Xiaoming Yang from the University of South Carolina for their help in the hemolysis study.

## Authors contributions

C.T. and A.W.D. were responsible for the conception, supervision, and financial support of the project. The work and the experiments were designed by C.T., A.W.D. and S. B. Transcriptomic analysis was conducted by J.L., A.B.C. and E.O. All other studies were conducted by S.B., A.A., M.W.H. and A.P. All data were analyzed by S.B., A.A., C.T. and A.W.D. The manuscript was written by S.B., C.T., and A.W.D. All authors approved the final version of the manuscript.

## Data availability

The main data supporting the results in this study are available within the article and its Supplementary Information. Transcriptomic datasets generated during the study are available online in the Gene Expression Omnibus database (https://www.ncbi.nlm.nih.gov/geo; Token No. erijcaekrzitnmb). The raw and analyzed datasets generated during the study are available for research purposes from the corresponding authors upon reasonable request.

## Competing interests

All authors declare no competing interests.

### Additional information

## Supplementary information

The online version contains supplementary material available at https://doi.org/10.1038/

**Correspondence and requests for materials** should be addressed to Chuanbing Tang or Alan W. Decho.

## Table of Contents Graphic

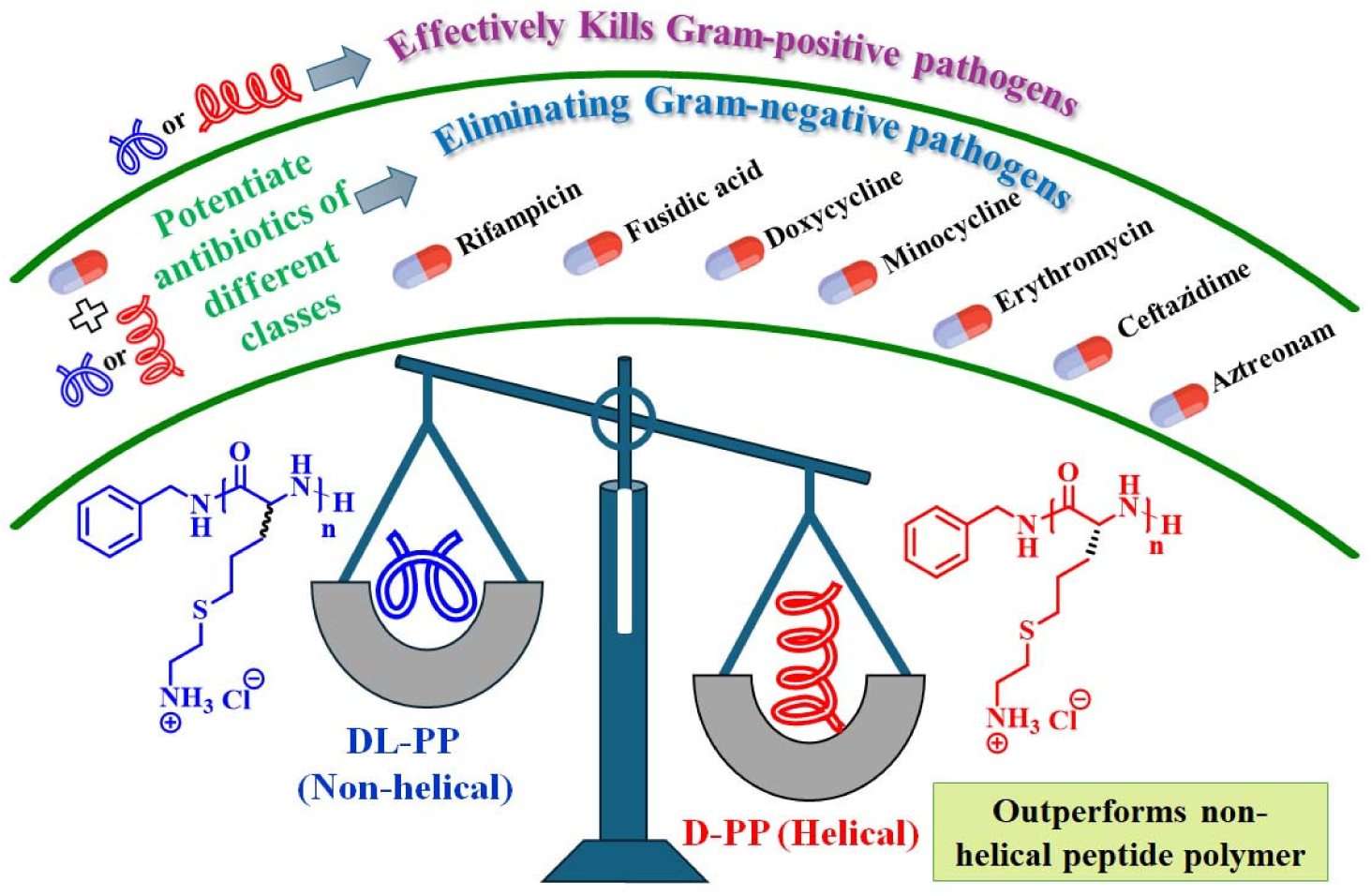

