## Supplemental for "The role of secondary structures of peptide polymers on antimicrobial efficacy and antibiotic potentiation"

#### Experimental Methods

**Material and instrumentations:** Reagents were purchased from commercial sources and used as received. D/DL-allylglycines and benzylamine were purchased from Ambeed Chemicals. Triphosgene, cysteamine hydrochloride and propylene oxide (PO) were purchased from TCI Chemicals. Hexafluoro isopropyl alcohol (HFIP) and antibiotics were purchased from Chem-Impex Chemicals. 2,2-Dimethoxy-2-phenylacetophenone (DMPA) and dyes for biological assays were bought from Mark Millipore. Common solvents such as hexane*, N*-dimethylformamide (DMF, 99.9%), dimethyl sulfoxide (DMSO) tetrahydrofuran (THF), methanol, etc. were brought from VWR International. Deuterated solvent, dimethyl sulfoxide DMSO-$d_6$ was purchased from Cambridge Isotope Laboratories. All bacterial strains including multidrug-resistant (MDR) strains were purchased from ATCC, USA. The general supplies for microbiological studies (e.g. bacterial media, buffers, well plates, centrifuge tubes etc.) were brought from VWR International.

Bruker Avance IV HD 400 spectrometer was employed to record proton and carbon Nuclear Magnetic Resonance (NMR) spectrum of monomers and polymers. In general, samples (5-8 mg) were dissolved in appropriate deuterated NMR solvents of ~600 µL before the NMR experiments. Gass chromatography-mass spectrometry (GC-MS) was used for the mass spectrometry of the monomers. Fourier Transform Infrared Spectrometry (FT-IR) spectra were recorded using a PerkinElmer spectrum 100 FTIR spectrometer incorporated with a diamond crystal plate as the reflector. Measurements for each sample were collected over a spectrum range of 4000–400 $cm^{-1}$. Circular dichroism (CD) spectroscopy was conducted using the Jasco J-815 Spectropolarimeter within the wavelength of 190-250 nm. Zeta potential measurements of the polymers were conducted using a Zetasizer Nano ZS (Malvern Instruments, Malvern, UK), equipped with a 4.0 mW 633 nm He-Ne laser and a detector positioned at 173 degrees. Each sample was prepared by dissolving the dry polymer in filtered deionized water with 25% DMSO. The typical concentration for all samples was 1.0 mg/mL. Data analysis was conducted using the general-purpose algorithms available in Zetasizer Software (version 7.11). Each sample measurement was carried out in triplicate, and results are presented as the average at a temperature of 25 °C. Optical density (OD) and fluorescence intensity measurements for the biological assays were conducted using a SpectraMax M5 Multimode Microplate Reader (Molecular Devices). For fluorescent imaging to assess LIVE/DEAD cells within the biofilm matrix, a Zeiss LSM 410 Confocal Laser Scanning Microscope (CLSM) was employed, providing high-resolution imaging into cell viability. Zeiss Gemini500 FE-SEM was employed to perform electron microscopy of the bacteria.

**Bactericidal kinetics against planktonic cells:**[1] Bacterial cells in the mid-log phase (approximately $10^8$ CFU/mL) were diluted to about $10^5$ CFU/mL in a 1:1 mixture of normal saline and cation-adjusted Mueller Hinton Broth. The cells were treated with standalone polymer, antibiotic, or their combinations. Specifically, 150 µL of this bacterial suspension was combined with 50 µL of either the standalone compound or the polymer-antibiotic combinations, followed by incubation at 37 °C. For the negative control, 50 µL of media devoid of polymers, antibiotics or their combinations was combined with 150 µL of bacterial suspension. At various time points (0, 2, 4, 6, 8, and 24 hours), 20 µL samples from each treatment concentration were 10-fold serially diluted in 1×PBS. From these dilutions, 20 µL aliquots were spot-plated on tryptic soya agar (TSA) plates and incubated for 24 hours at 37 °C. After incubation, viable colonies were counted on the plates, with a detection limit of 50 CFU/mL for this experiment. All experiments were performed in duplicate to ensure reliability.

**Visual turbidity assay.**[2] Briefly, 5 mL of ~$10^5$ CFU/mL mid-log phase bacterial solution was treated individually with an aqueous solution of polymer, antibiotic and their combination at different concentrations for different bacteria. Then the compound-treated bacterial suspension was incubated at 37 °C under shaking conditions for 16-18 h. Afterwards, photographic images were captured for each experimental tube. Finally, 20 µL of bacterial suspension from each experimental tube was spread on the TSA plate and the plate was again incubated for 24 h at 37 °C under stationary conditions. In the end, photographic images of the agar plates were captured to visualize the bacterial growth or no growth.

**Bactericidal kinetics against dormant bacteria.** [1] A mid-log phase (6 h grown culture) bacterial culture was diluted to a 1:1000 ratio in nutrient broth (NB) and incubated at 37 ˚C for 16 h under shaking at 190 rpm to achieve stationary phase cells. Afterwards, the bacterial suspension was centrifuged (3500 rpm, 5 min) and resuspended in 1×PBS (pH = 7.4). Finally, 150 µL of the stationary phase bacteria (~5× $10^5$ CFU/mL) in 1×PBS was added to 50 µL of polymer, antibiotics and their combination with different concentrations in the same buffer. Similarly, polymyxin-B (16 µg/mL) was used as a control antibiotic for this study. The same volume of 1×PBS without any compound was considered untreated control. After 0, 4, 8 and 24 hours, 20 µL aliquots from that solution were serially diluted 10-fold in sterile saline. Then 20 µL solution from each dilution was spotted plated on TSA plates and after 24 h of incubation at 37 ˚C, the number of bacterial colonies was counted. The results were presented in a bar plot on a logarithmic scale, i.e. Log (CFU/mL) vs time points.

**Hemolysis:**[3] Blood was collected from mice in heparinized tubes and washed with 1xPBS +10% FBS. The peptide polymer solution (50 µL), prepared in 1xPBS at different concentrations, was added to the red blood cells (RBCs) in 96-well plates (150 µL). 1xPBS, with 0.2% Triton-X100 (TRX) served as maximum lysis control. The samples were incubated at 37°C for 1 h followed by centrifugation at 1500 rpm for 10 minutes. Supernatants (100 µL) were transferred to a 96-well plate and OD was measured at 450 nm. 1xPBS+10%FBS was used for background reading. The percentage of hemolysis was determined by using the following formula:

$$(OD_{tret} - OD_{background}) / (OD_{TRX\text{-}tret} - OD_{background}) \times 100$$

where $OD_{tret}$ corresponds to the absorbance of the compound-treated well, $OD_{background}$ stands for the absorbance of the negative controls (without compound), and $OD_{TRX\text{-}tret}$ is the absorbance of the triton-X-100 treated well. Each concentration had triplicate values and the $HC_{50}$ was determined by considering the average of triplicate O.D.

**Zeta potential of bacteria:**[4] Approximately $10^8$ CFU/mL of mid-log phase bacteria (*K. pneumonia* ATCC BAA 2473) was suspended in 1 mL sterile water. Afterwards, the fresh bacterial suspension was placed in a quartz cuvette (path length: 3 mm) and its zeta potential was measured immediately using Zetasizer Nano ZS in triplicates. Following this, the 10 μL aqueous solution of the peptide polymers of a particular concentration was added to the bacterial suspension and the change in zeta potential was further recorded. The whole experiment was separately performed for different polymers with different concentrations.

**Scanning electron microscopy (SEM):**[3] Briefly, 1 mL of ~$10^6$ CFU/mL mild log phase cells of MRSA ATCC BAA 44 were treated with polymer (8 μ/mL) in 1:1 mixture of Muller Hinton Broth (MHB) and normal saline in a 12 well plate. In the case of control, bacterial cells were not treated with the compound. Further, two sterilized coverslips (tissue culture treated and 13 mm diameter) were added into each 1mL bacterial suspension (in 12 wells) immediately in case of treated and untreated samples and incubated for 16 h at 37 °C under static conditions. Afterwards, the coverslips were taken out from the wells and processed for SEM imaging. First, the coverslips (bearing the bacterial cells) were incubated with 2.5% glutaraldehyde solution in 0.1 M sodium cacodylate buffer for 2 hours at 4 °C. Next, the same coverslips were incubated in a 1% aqueous solution of osmium tetroxide for another hour at 4 °C and they were further washed with 0.1 M sodium cacodylate buffer to remove trace amount of toxic osmium tetroxide. Next, the coverslips were gradually incubated with 50%, 70%, 80%, 95% and 100% ethanol and each incubation step was for 10 min at room temperature. Finally, the coverslips (containing fixed and dehydrated bacterial cells) were dried using the critical point chamber, then coated with gold using a Denton Desk II sputter coater for 60 seconds, and images were captured using Zeiss Gemini500 FE-SEM. The images were further processed with Fiji (ImageJ-win64).

**Resistance study:**[2,5] The assay was performed for D-PP, doxycycline as well as the combined treatment of D-PP and doxycycline against *P. aeruginosa* ATCC BAA 2108. In the case of combination, the MIC value of doxycycline was evaluated by keeping the concentration of D-PP constant at 4 μg/mL. To compare the result, Gram-negative antibiotic, polymyxin-B was incorporated in the study. By following the protocol described in the antibacterial assay section, the MIC values of D-PP, polymyxin-B and doxycycline (in the presence of 4 μg/mL of D-PP) were determined against the same *P. aeruginosa*. The next MIC assay was performed, where the bacterial suspension was then prepared from the bacteria grown at the sub-MIC of compound concentration of the first-day experiment. By following the same protocol, 14 subsequent passages were repeated. To evaluate the propensity for resistance development, the fold increase in MIC values was plotted against the passage number. MIC values were calculated based on visual observations of optical density (OD) readings obtained from triplicate experiments. The fold increase in MIC was determined by dividing the initial MIC value (corresponding to the first passage) by the MIC values recorded at each subsequent passage.

### Supporting Figures

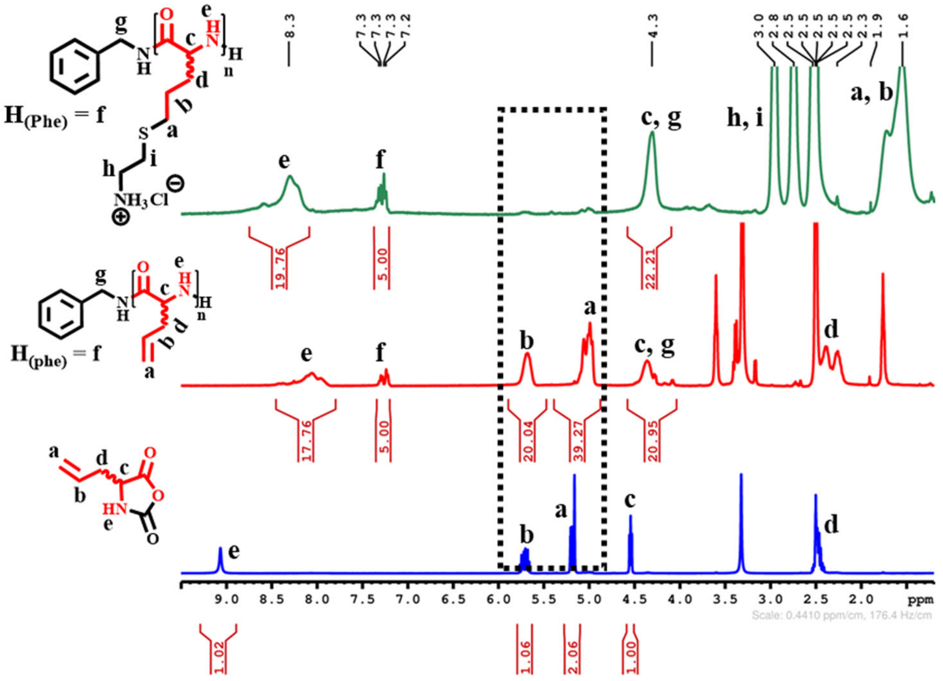

**Figure S1.** Stacked ¹H-NMR spectra of DL-allylglycine NCA, corresponding poly(DL-allylglycine) and its post-functionalized cationic DL-peptide polymer (DL-PP).

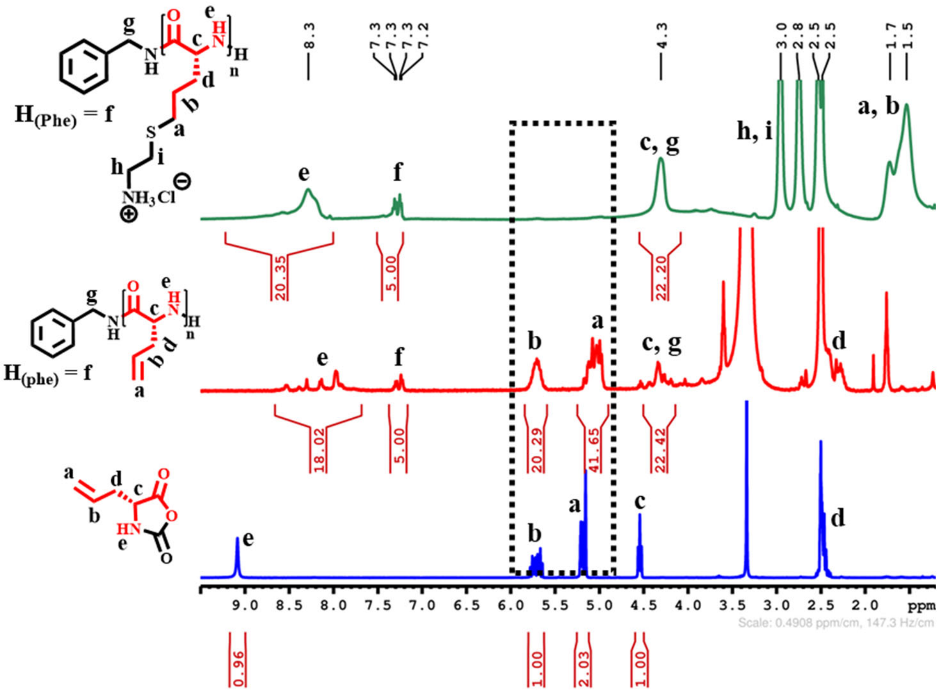

**Figure S2**. Stacked ¹H-NMR spectra of D-allylglycine NCA, corresponding poly(D-allylglycine) and its post-functionalized cationic D-peptide polymer (D-PP).

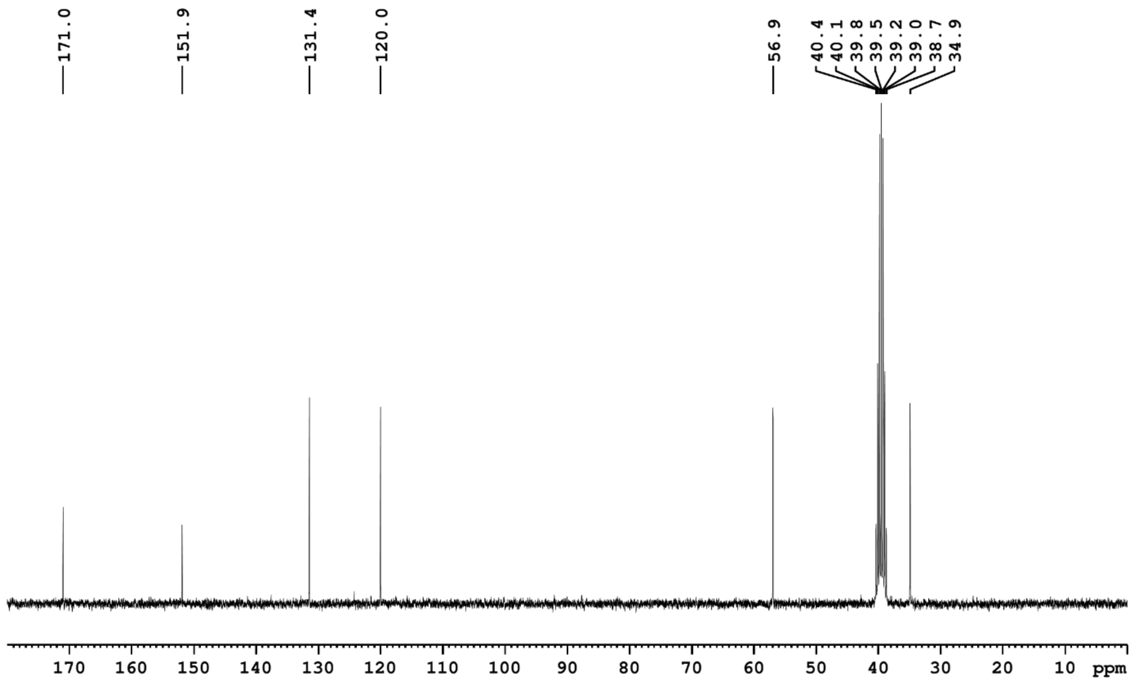


**Figure S3.** The [13]C-NMR spectrum of DL-allylglycine NCA. The NMR was recorded in DMSO-$d_6$, and the solvent peak was calibrated at a chemical shift of 39.52 ppm.

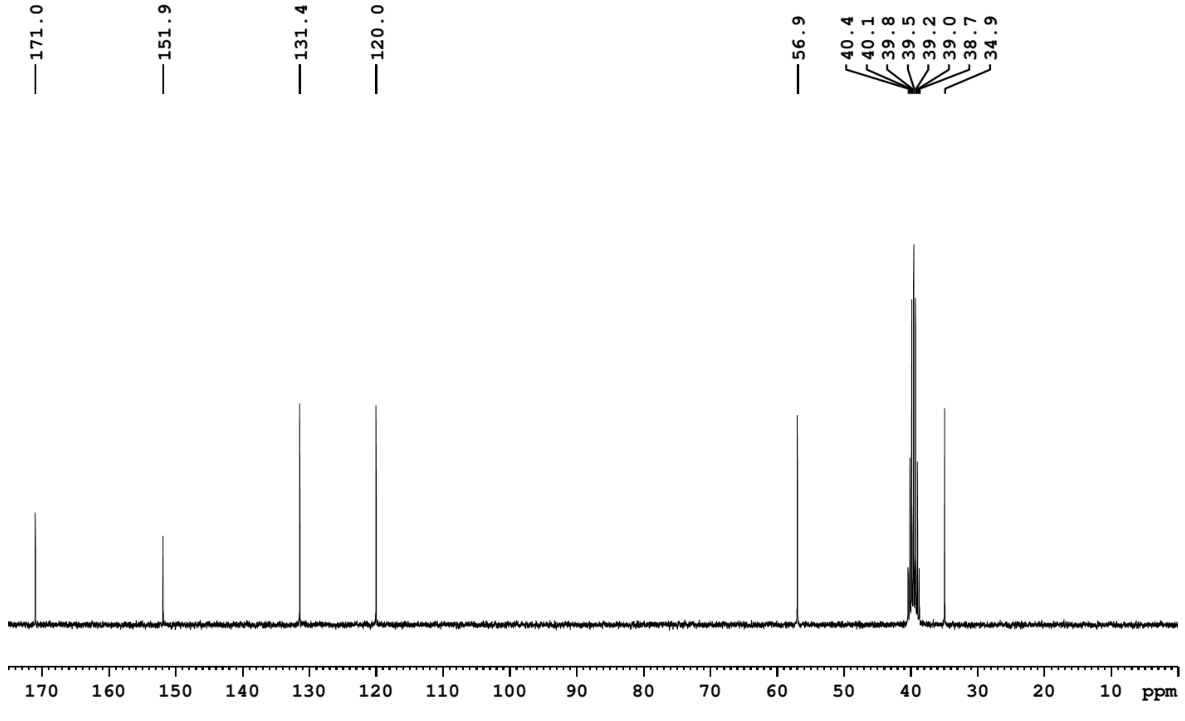


**Figure S4.** The [13]C-NMR spectrum of D-allylglycine NCA. The NMR was recorded in DMSO-$d_6$, and the solvent peak was calibrated at a chemical shift of 39.52 ppm.

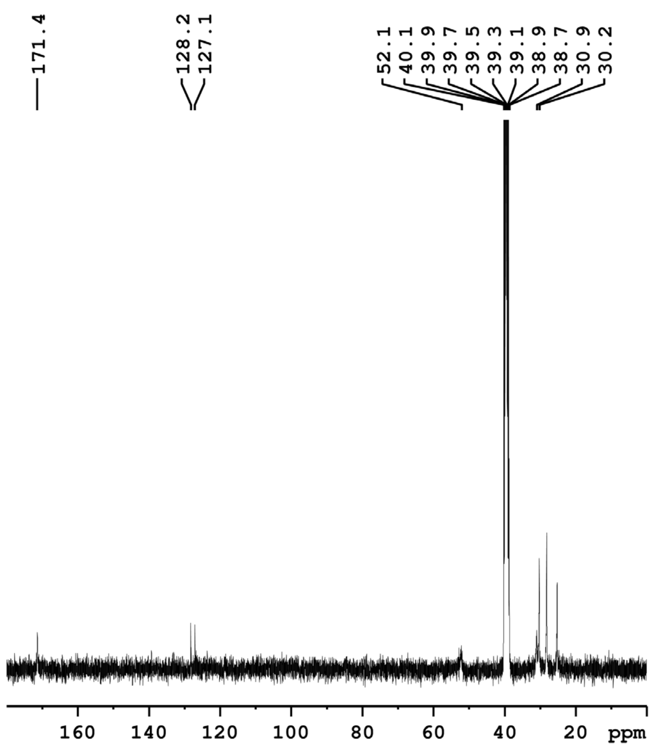


**Figure S5.** The $^{13}$C-NMR spectrum of DL-PP. The NMR was recorded in DMSO- $d_6$, and the solvent peak was calibrated at a chemical shift of 39.52 ppm.

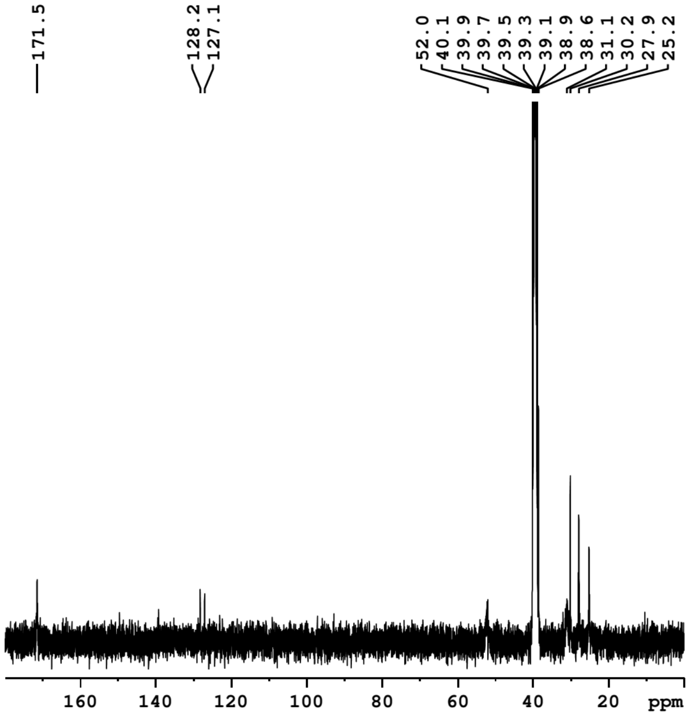


**Figure S6.** The $^{13}$C-NMR spectrum of D-PP. The NMR was recorded in DMSO-$d_6$, and the solvent peak was calibrated at a chemical shift of 39.52 ppm.

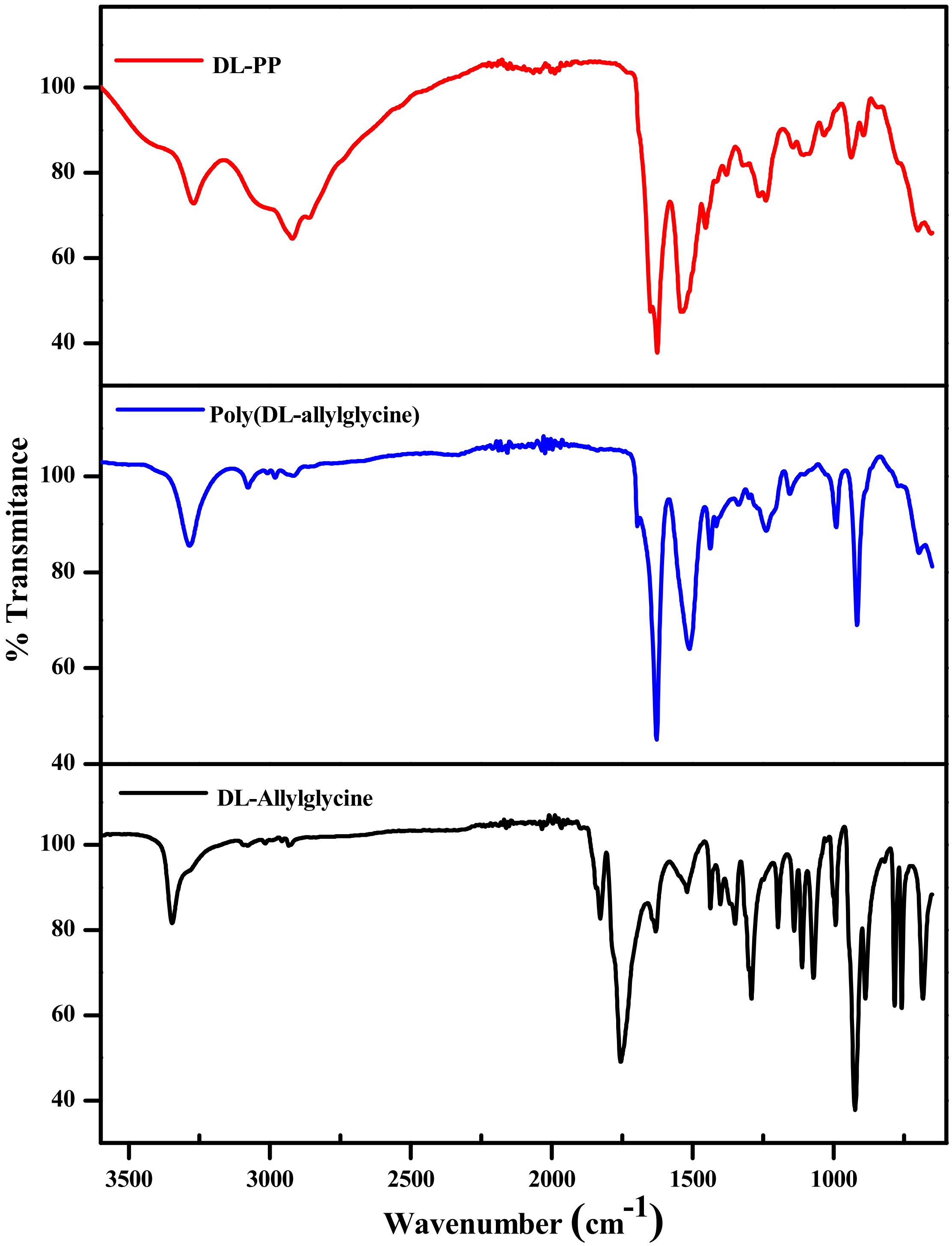


**Figure S7.** The FT-IR spectra of DL-PP, its precursor polymer, poly(DL-allylglycine) and monomer, DL-allylglycine NCA.

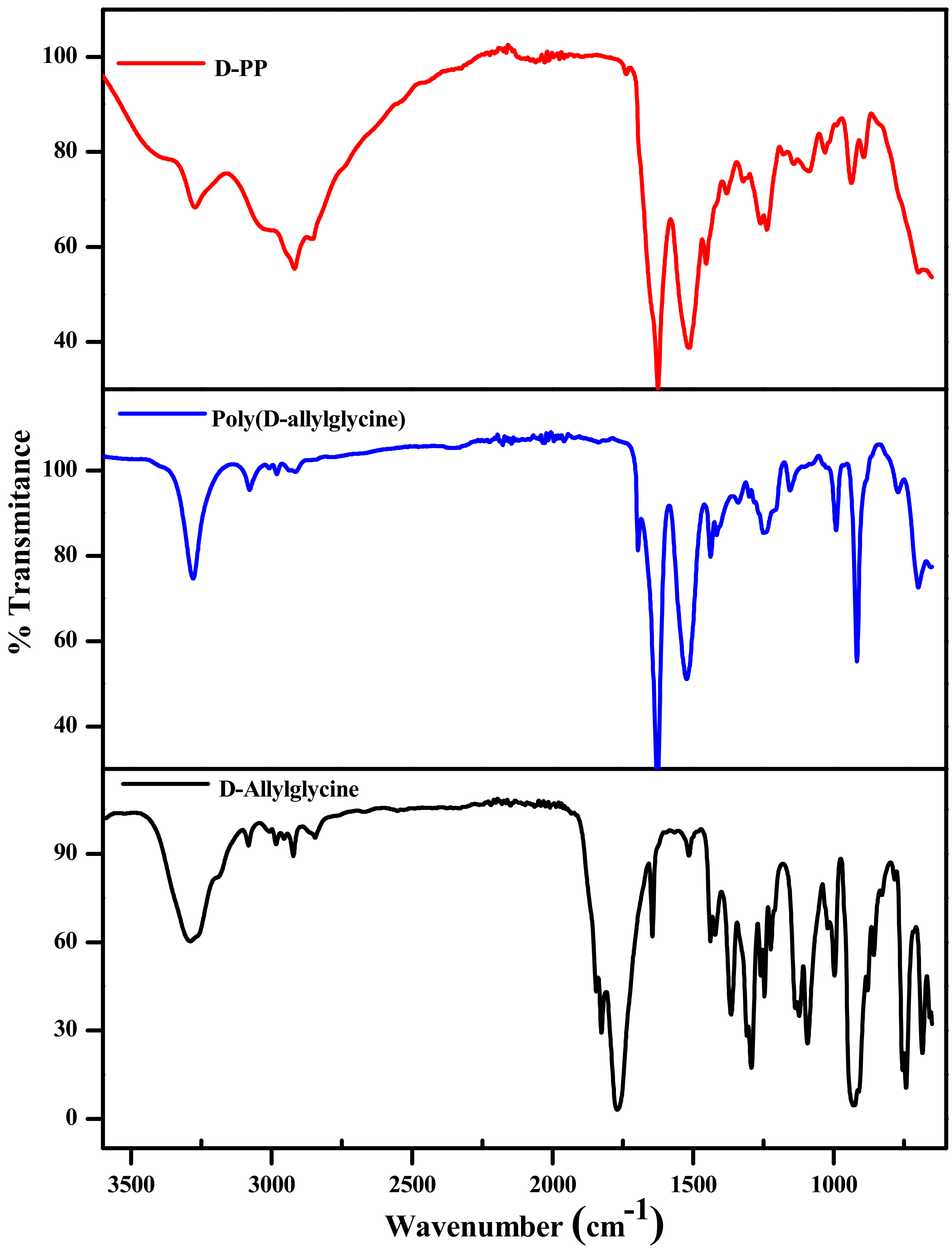


**Figure S8.** The FT-IR spectra of D-PP, its precursor polymer, poly(D-allylglycine) and monomer, D-allylglycine NCA.

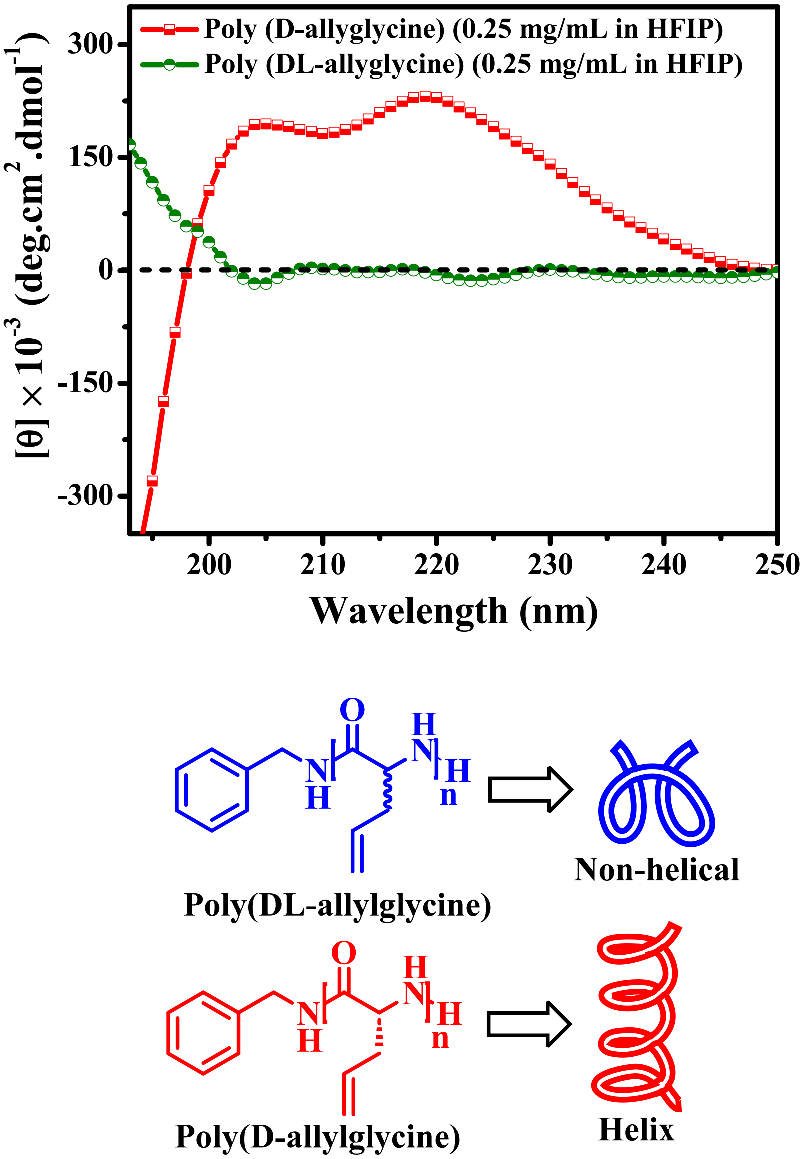


**Figure S9.** The precursor D-polymer adopted a secondary structure in biomimicking solvent, HFIP. Circular dichroism (CD) spectra of (a) poly(DL-allylglycine) and poly(D-allylglycine) in HFIP.

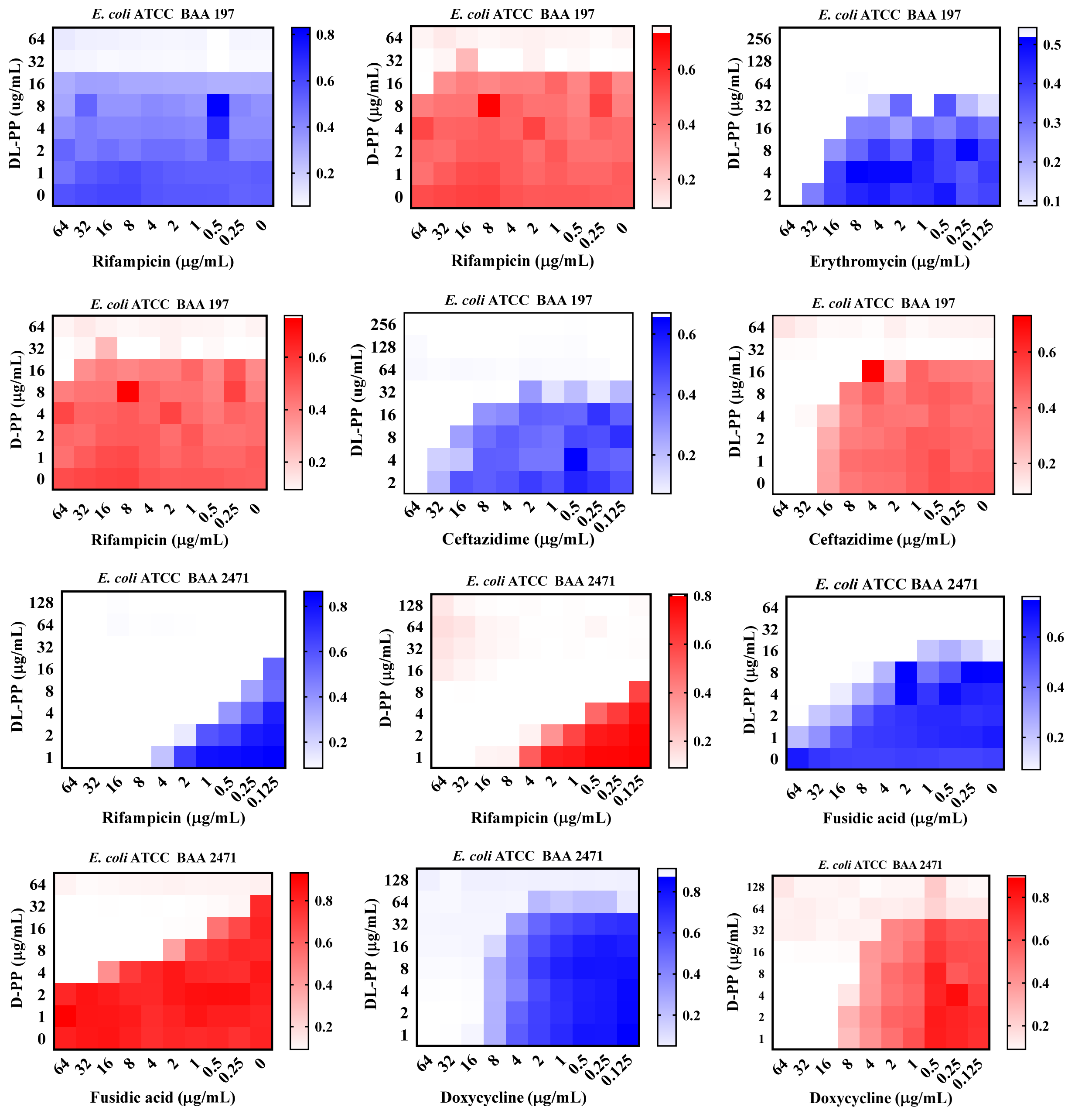


**Figure S10.** Heatmap represents the checkerboard assays displaying the potentiation of antibiotics of different classes against MDR *E. coli* ATCC BAA 197 and *E. coli* ATCC BAA 2471 by helical (D-PP) and non-helical peptide polymers (DL-PP) at variable concentrations. The decrease in red or blue colour intensity represents the decrease in microbial cell density.

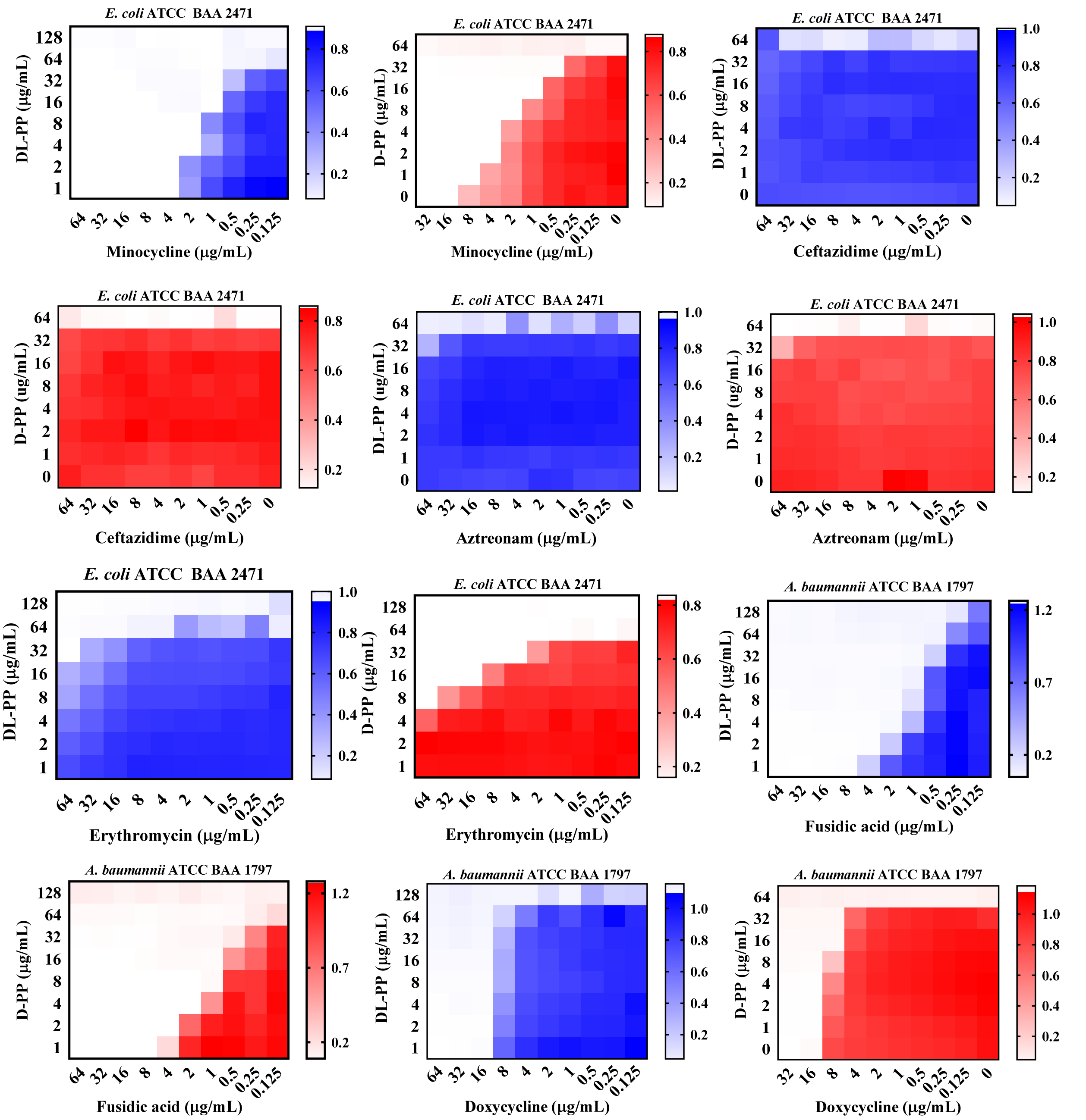


**Figure S11.** Heatmap represents the chequerboard assays displaying the potentiation of antibiotics of different classes against MDR *E. coli* ATCC BAA 2471 and *A. baumannii* ATCC BAA 1797 by helical (D-PP) and non-helical peptide polymers (DL-PP) at variable concentrations. The decrease in red or blue colour intensity represents the decrease in microbial cell density.

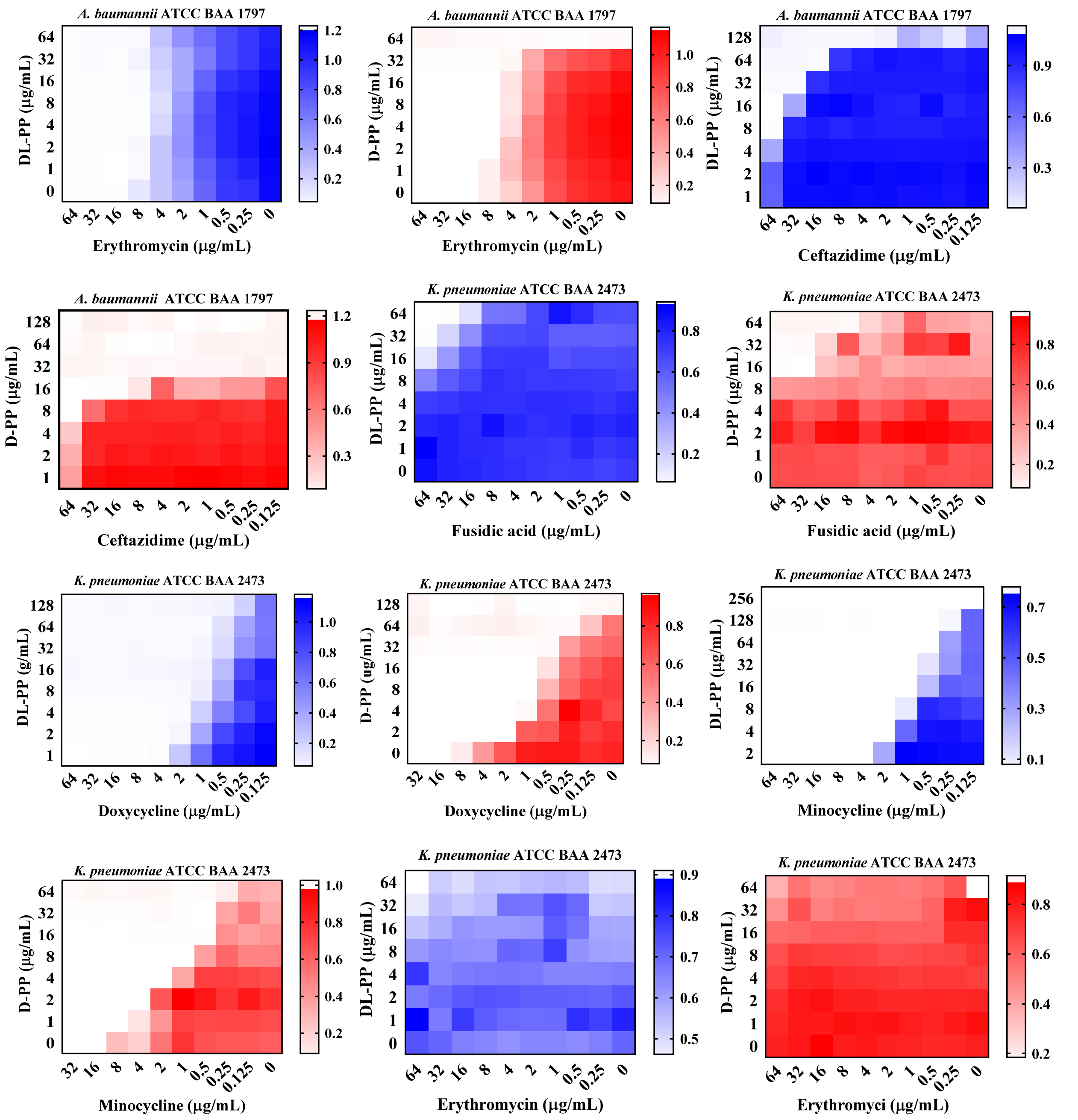


**Figure S12.** Heatmap represents the checkerboard assays displaying the potentiation of antibiotics of different classes against MDR *A. baumannii* ATCC BAA 1797 and *K. pneumoniae* ATCC BAA 2473 by helical (D-PP) and non-helical peptide polymers (DL-PP) at variable concentrations. The decrease in red or blue colour intensity represents the decrease in microbial cell density.

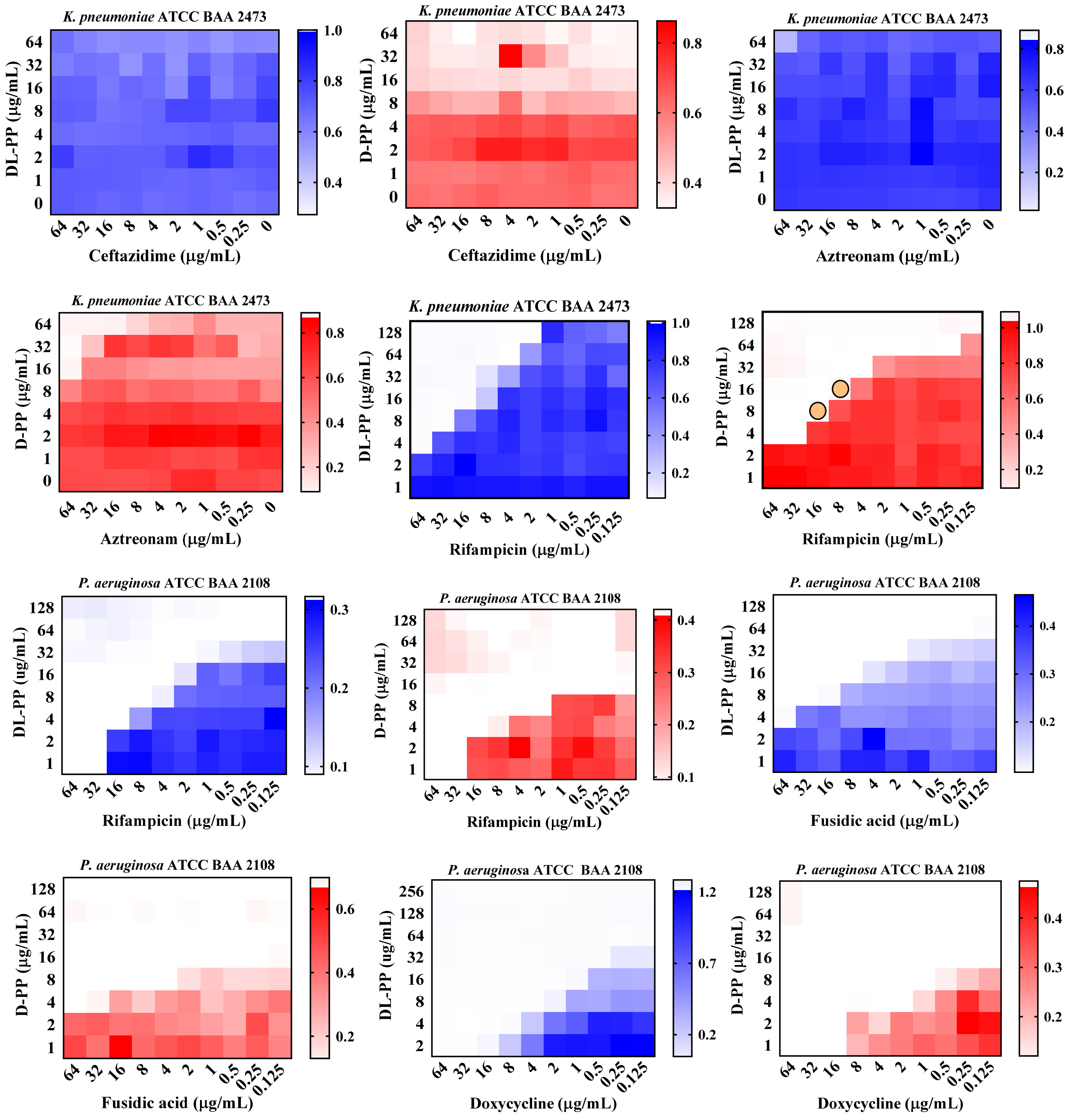


**Figure S13.** Heatmap represents the checkerboard assays displaying the potentiation of antibiotics of different classes against MDR *K. pneumoniae* ATCC BAA 2473 and *P. aeruginosa* ATCC BAA 2108 by helical (D-PP) and non-helical peptide polymers (DL-PP) at variable concentrations. The decrease in red or blue colour intensity represents the decrease in microbial cell density.

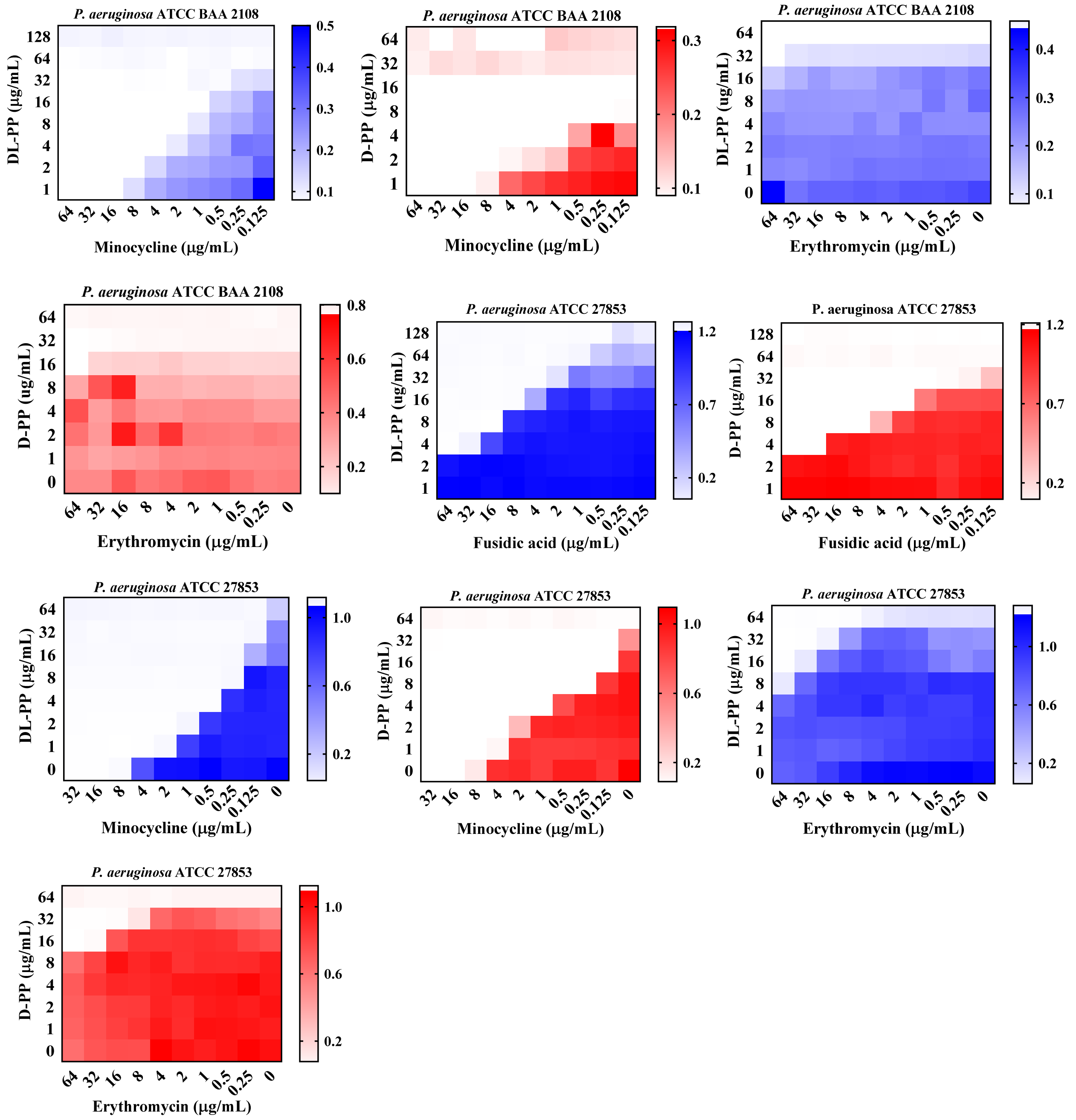


**Figure S14.** Heatmap represents the checkerboard assays displaying the potentiation of antibiotics of different classes against MDR *P. aeruginosa* ATCC BAA 2108 and *P. aeruginosa* ATCC 27853 by helical (D-PP) and non-helical peptide polymers (DL-PP) at variable concentrations. The decrease in red or blue colour intensity represents the decrease in microbial cell density.

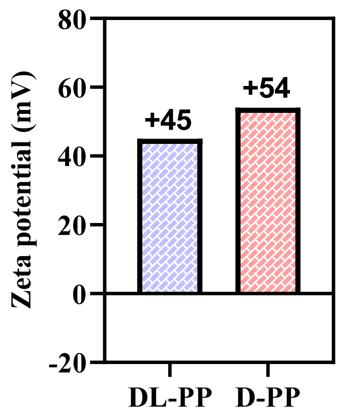


**Figure S15**. Zeta potential of aqueous solution of peptide polymers (D-PP and DL-PP). The concentration of the polymers was 1 mg/mL.

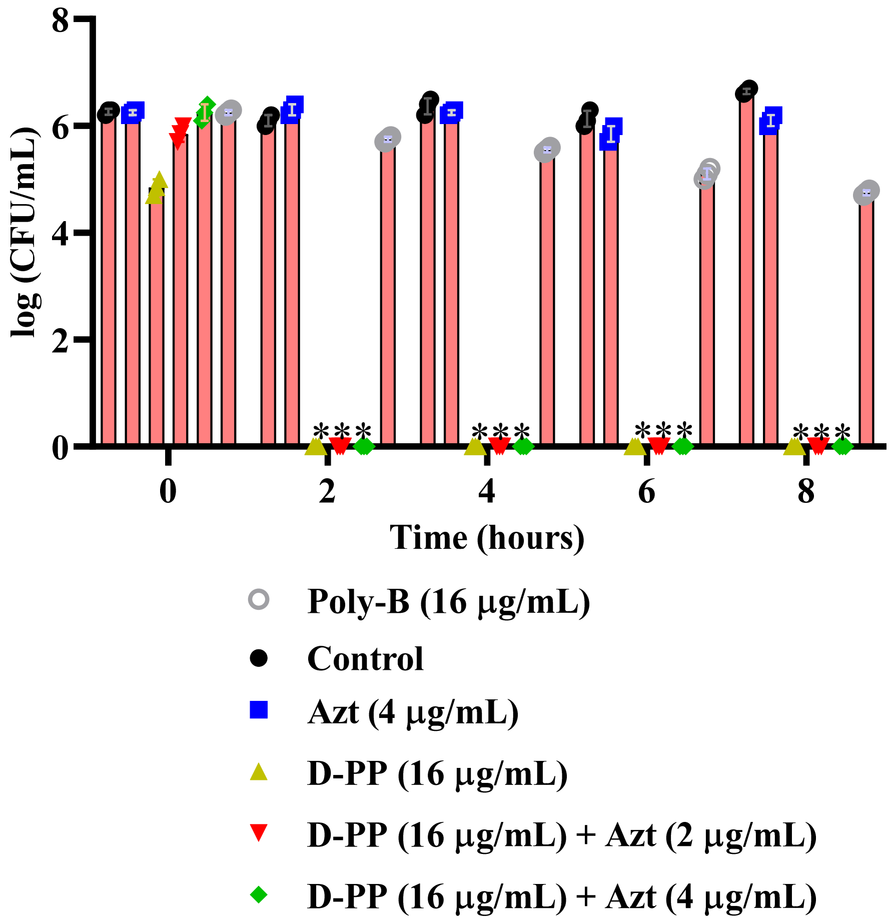


**Figure S16.** Efficacy of D-PP and aztreonam (Azt) combinations against dormant MDR A. *baumannii* ATCC BAA 1797 (Stationary cells). Arteries indicate <50 CFU/mL.

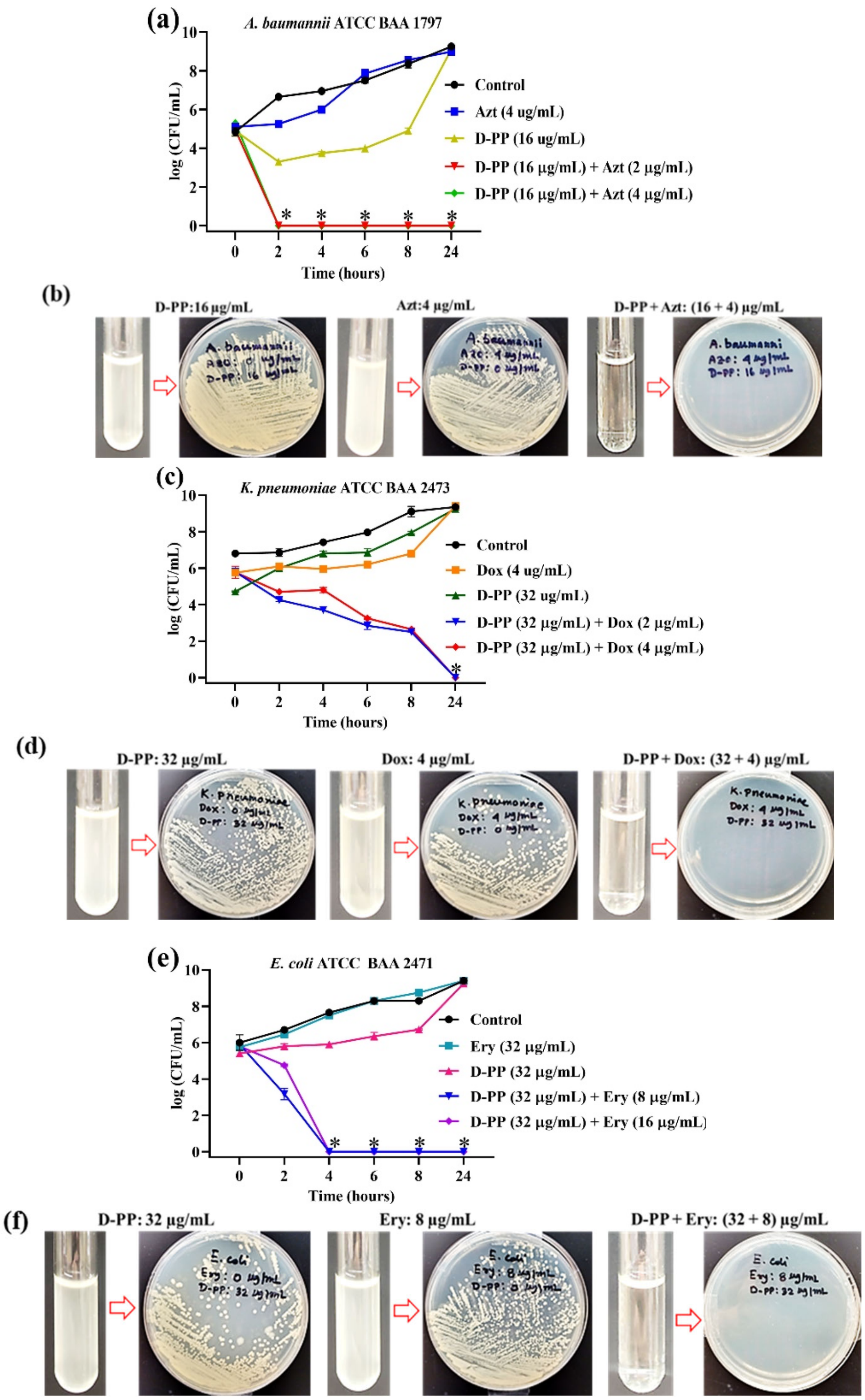


**Figure S17. D-PP is bactericidal in nature.** (a) Bactericidal killing kinetic and (b) visual turbidity of D-PP and aztreonam (Azt) combination, (c) Bactericidal killing kinetic and (d) visual turbidity of D-PP and doxycycline (Dox) combination, (e) Bactericidal killing kinetic and (f) visual turbidity of D-PP and erythromycin (Ery) combination. Visual turbidity, combined with plating treated and untreated bacterial suspensions on agar, offers a straightforward representation of bactericidal kinetics. Asterisk represents <50 CFU/mL.

**The helical polymer adjuvant-antibiotic combinations exhibited potent bactericidal activity against both planktonic bacteria.** To evaluate the rate of bacterial eradication, we performed killing kinetics on three representative bacterial species using polymer-antibiotic combinations (Figure S17). Turbidity measurements (reflecting bacterial presence) and plating assays of treated versus untreated bacterial suspensions confirmed the bactericidal effects. For example, a combination of D-PP (16 µg/mL) and aztreonam (2–4 µg/mL) achieved rapid bactericidal action, reducing *A. baumannii* planktonic cells by ~4.8 logs within 2 hours (Figure S17a). No turbidity or bacterial colonies were observed in the media or on solid agar for samples treated with D-PP and aztreonam, indicating complete bacterial elimination. In contrast, treatments with D-PP or aztreonam alone resulted in high bacterial viability, evident turbidity, and bacterial colonies on agar (Figure S17b). Similarly, we observed a complete killing of MDR *K. pneumoniae* with ~6 log reduction within 24 h when treated with the combination of D-PP (32 µg/mL) and doxycycline (2 and 4 µg/mL) unlike the standalone treatments (Figure S17c). There was no visual turbidity in growth media and no colonies on the solid agar for *K. pneumoniae* exposed to the doxycycline (32 µg/mL)-D-PP (4 µg/mL) combination (Figure S17d). In another case, the combination of D-PP (32 µg/mL) with erythromycin (8–16 µg/mL) fully eradicated MDR *E. coli* within 4 hours, yielding a ~6 log reduction, while single treatments with either D-PP or erythromycin resulted in a 9-log bacterial increase within 24 hours, comparable to untreated controls (Figure S17e). The erythromycin-D-PP combination produced clear media and agar with no detectable bacterial colonies (Figure S17f). These findings highlighted the swift and potent bactericidal effects of D-PP-antibiotic combinations, significantly surpassing individual treatments.

**Table S1.** Antibacterial efficacy of the combination of D-PP or DL-PP and antibiotics of different classes against the tested multidrug-resistant Gram-negative bacteria.

| Bacteria and antibiotics | MIC of antibiotic (-D/DL-PP) | MIC of DL-PP | MIC of D-PP | MIC of antibiotic (+DL-PP) | MIC of antibiotic (+D-PP) |
|---|---|---|---|---|---|
| **Rifampicin** | | | | | |
| *E. coli* ATCC BAA 197 | >1024 | 32-64 | 32 | >64 (+16) | 64 (+16) |
| *E. coli* ATCC BAA 2471 | 16 | 256 | 64 | 0.5 (+8) | 0.25 (+8) |
| *A. baumannii* ATCC BAA 1797 | 1 | 256 | 64 | N.D.* | N.D. |
| *K. pneumoniae* ATCC BAA 2473 | >256 | 256 | 64 | 16 (+16) | 8 (+16) |
| *P. aeruginosa* ATCC BAA 2108 | 32 | 64 | 16 | 16 (+4) | 8 (+4) |
| *P. aeruginosa* ATCC 27853 | 32 | 256 | 64 | 0.5 (+16) | 0.25 (+16) |
| **Fusidic acid** | | | | | |
| *E. coli* ATCC BAA 197 | >256 | 32-64 | 32 | 8 (+8) | 16 (+8) |
| *E. coli* ATCC BAA 2471 | >256 | 256 | 64 | 2 (+16) | 1 (+16) |
| *A. baumannii* ATCC BAA 1797 | 128 | 256 | 64 | 2 (+8) | 1 (+8) |
| *K. pneumoniae* ATCC BAA 2473 | >256 | 256 | 64 | >64 (+16) | 32 (+16) |
| *P. aeruginosa* ATCC BAA 2108 | >256 | 64 | 16 | 16 (+8) | 4 (+8) |
| *P. aeruginosa* ATCC 27853 | >256 | 256 | 64 | 8 (+16) | 2 (+16) |
| **Doxycycline** | | | | | |
| *E. coli* ATCC BAA 197 | <1 | 32-64 | 32 | N.D. | N.D. |
| *E. coli* ATCC BAA 2471 | 64 | 256 | 64 | 16 (+8) | 8 (+8) |
| *A. baumannii* ATCC BAA 1797 | 16 | 256 | 64 | 16 (+16) | 8 (+16) |
| *K. pneumoniae* ATCC BAA 2473 | 16 | 256 | 64 | 2 (+16) | 1 (+16) |
| *P. aeruginosa* ATCC BAA 2108 | 32 | 64 | 16 | 8 (+4) | 2 (+4) |
| *P. aeruginosa* ATCC 27853 | 16 | 256 | 64 | 0.25 (+16) | 0.12 (+16) |
| **Minocycline** | | | | | |
| *E. coli* ATCC BAA 197 | <1 | 32-64 | 32 | N.D. | N.D. |
| *E. coli* ATCC BAA 2471 | 16 | 256 | 64 | 1 (+16) | 1 (+16) |
| *A. baumannii* ATCC BAA 1797 | 1 | 256 | 64 | N.D. | N.D. |
| *K. pneumoniae* ATCC BAA 2473 | 16 | 256 | 64 | 1 (+16) | 0.5 (+16) |
| *P. aeruginosa* ATCC BAA 2108 | 32 | 64 | 16 | 4 (+4) | 1 (+4) |
| *P. aeruginosa* ATCC 27853 | 16 | 256 | 64 | 0.25 (+16) | 0.12 (+16) |
| **Erythromycin** | | | | | |
| *E. coli* ATCC BAA 197 | 128 | 32-64 | 32 | 32 (+4) | 32 (+4) |
| *E. coli* ATCC BAA 2471 | 1024 | 256 | 64 | >64 (+16) | 16 (+16) |
| *A. baumannii* ATCC BAA 1797 | 16 | 256 | 64 | 8 (+4) | 8 (+4) |
| *K. pneumoniae* ATCC BAA 2473 | >256 | 256 | 64 | >64 (+16) | >64 (+16) |
| *P. aeruginosa* ATCC BAA 2108 | 256 | 64 | 16 | >64 (+4) | >64 (+4) |
| *P. aeruginosa* ATCC 27853 | 256 | 256 | 64 | 64 (+16) | 32 (+16) |
| **Ceftazidime** | | | | | |
| *E. coli* ATCC BAA 197 | 32-64 | 32-64 | 32 | 16 (+16) | 8 (+16) |
| *E. coli* ATCC BAA 2471 | >1024 | 256 | 64 | >64 (+16) | >64 (+16) |
| *A. baumannii* ATCC BAA 1797 | 128 | 256 | 64 | 64 (+16) | 16 (+16) |
| *K. pneumoniae* ATCC BAA 2473 | >1024 | 256 | 64 | >64 (+16) | >64 (+16) |
| *P. aeruginosa* ATCC BAA 2108 | 4 | 64 | 16 | N.D. | N.D. |
| *P. aeruginosa* ATCC 27853 | 2 | 256 | 64 | N.D. | N.D. |
| **Aztreonam** | | | | | |
| *E. coli* ATCC BAA 197 | 2 | 32-64 | 32 | N.D. | N.D. |
| *E. coli* ATCC BAA 2471 | >1024 | 256 | 64 | >64 (+16) | >64 (+16) |
| *A. baumannii* ATCC BAA 1797 | 64 | 256 | 64 | 8 (+16) | 2 (+16) |
| *K. pneumoniae* ATCC BAA 2473 | >1024 | 256 | 64 | >64 (+16) | 64 (+16) |
| *P. aeruginosa* ATCC BAA 2108 | 2 | 64 | 16 | N.D. | N.D. |
| *P. aeruginosa* ATCC 27853 | 4 | 256 | 64 | N.D. | N.D. |

*N.D. refers to “not determined”

**Table S2.** Summary of PF, FIC of adjuvants, FICI of the combination assays against the tested multidrug-resistant Gram-negative bacteria.

| Bacteria and antibiotics | PF (DL-PP) | PF (D-PP) | FIC (DL-PP) | FIC (D-PP) | FICI (DL-PP) | FICI (D-PP) |
|---|---|---|---|---|---|---|
| **Rifampicin** | | | | | | |
| *E. coli* ATCC BAA 197 | N.D. | >16 | N.D.* | 16 | N.D. | <0.56 (Add) |
| *E. coli* ATCC BAA 2471 | 32 | 64 | 8 | 8 | 0.06 (Syn) | 0.14 (Syn) |
| *A. baumannii* ATCC BAA 1797 | N.D. | N.D. | N.D. | N.D. | N.D. | N.D. |
| *K. pneumoniae* ATCC BAA 2473 | 16 | 32 | 16 | 16 | 0.12 (Syn) | 0.28 (Syn) |
| *P. aeruginosa* ATCC BAA 2108 | 2 | 4 | 4 | 4 | 0.56 (Add) | 0.5 (Syn) |
| *P. aeruginosa* ATCC 27853 | 64 | 128 | 16 | 16 | 0.08 (Syn) | 0.25 (syn) |
| **Fusidic acid** | | | | | | |
| *E. coli* ATCC BAA 197 | >32 | >16 | 8 | 8 | <0.28 (Syn) | <0.31 (Syn) |
| *E. coli* ATCC BAA 2471 | >128 | >256 | 16 | 16 | <0.07 (Syn) | <0.25 (Syn) |
| *A. baumannii* ATCC BAA 1797 | 64 | 128 | 8 | 8 | 0.04 (Syn) | 0.13 (Syn) |
| *K. pneumoniae* ATCC BAA 2473 | ≥1 | 8 | 16 | 16 | ≤1.06 (Syn) | <0.37 (Syn) |
| *P. aeruginosa* ATCC BAA 2108 | 16 | 64 | 8 | 8 | 0.31 (Syn) | 0.51 (Add) |
| *P. aeruginosa* ATCC 27853 | 32 | 128 | 16 | 16 | 0.09 (Syn) | 0.26 (Syn) |
| **Doxycycline** | | | | | | |
| *E. coli* ATCC BAA 197 | N.D. | N.D. | N.D. | N.D. | N.D. | N.D. |
| *E. coli* ATCC BAA 2471 | 4 | 8 | 8 | 8 | 0.25 (Syn) | 0.25 (Syn) |
| *A. baumannii* ATCC BAA 1797 | 1 | 2 | 16 | 16 | 1.06 (Ind) | 0.75 (Add) |
| *K. pneumoniae* ATCC BAA 2473 | 8 | 16 | 16 | 16 | 0.18(Syn) | 0.31 (Syn) |
| *P. aeruginosa* ATCC BAA 2108 | 4 | 16 | 4 | 4 | 0.26 (Syn) | 0.12 (Syn) |
| *P. aeruginosa* ATCC 27853 | 64 | 128 | 16 | 16 | 0.08 (Syn) | 0.25 (Syn) |
| **Minocycline** | | | | | | |
| *E. coli* ATCC BAA 197 | N.D. | N.D. | N.D. | N.D. | N.D. | N.D. |
| *E. coli* ATCC BAA 2471 | 16 | 16 | 16 | 16 | 0.12 (Syn) | 0.31 (Syn) |
| *A. baumannii* ATCC BAA 1797 | N.D. | N.D. | N.D. | N.D. | N.D. | N.D. |
| *K. pneumoniae* ATCC BAA 2473 | 16 | 32 | 16 | 16 | 0.12 (Syn) | 0.28 (Syn) |
| *P. aeruginosa* ATCC BAA 2108 | 8 | 32 | 4 | 4 | 0.18 (Syn) | 0.28 (Syn) |
| *P. aeruginosa* ATCC 27853 | 64 | 128 | 16 | 16 | 0.08 (Syn) | 0.26 (Syn) |
| **Erythromycin** | | | | | | |
| *E. coli* ATCC BAA 197 | 4 | 4 | 4 | 4 | 0.37 (Syn) | 0.37 (Syn) |
| *E. coli* ATCC BAA 2471 | N.D. | 64 | 16 | 16 | N.D. | 0.26 (Syn) |
| *A. baumannii* ATCC BAA 1797 | 2 | 2 | 4 | 4 | 0.51 (Add) | 0.56 (Add) |
| *K. pneumoniae* ATCC BAA 2473 | N.D. | N.D. | N.D. | N.D. | N.D. | N.D. |
| *P. aeruginosa* ATCC BAA 2108 | N.D. | N.D. | N.D. | N.D. | N.D. | N.D. |
| *P. aeruginosa* ATCC 27853 | 4 | 8 | 16 | 16 | 0.31 (Syn) | 0.37 (Syn) |
| **Ceftazidime** | | | | | | |
| *E. coli* ATCC BAA 197 | 2-4 | 4-8 | 16 | 16 | 0.5-1 (Syn-Add) | 0.6-0.7 (Add) |
| *E. coli* ATCC BAA 2471 | N.D. | N.D. | N.D. | N.D. | N.D. | N.D. |
| *A. baumannii* ATCC BAA 1797 | 2 | 8 | 16 | 16 | 0.56 (Add) | 0.37 (Syn) |
| *K. pneumoniae* ATCC BAA 2473 | N.D. | N.D. | N.D. | N.D. | N.D. | N.D. |
| *P. aeruginosa* ATCC BAA 2108 | N.D. | N.D. | N.D. | N.D. | N.D. | N.D. |
| *P. aeruginosa* ATCC 27853 | N.D. | N.D. | N.D. | N.D. | N.D. | N.D. |
| **Aztreonam** | | | | | | |
| *E. coli* ATCC BAA 197 | N.D. | N.D. | N.D. | N.D. | N.D. | N.D. |
| *E. coli* ATCC BAA 2471 | N.D. | N.D. | N.D. | N.D. | N.D. | N.D. |
| *A. baumannii* ATCC BAA 1797 | 8 | 32 | 16 | 16 | 0.18 (Syn) | 0.28 (Syn) |
| *K. pneumoniae* ATCC BAA 2473 | N.D. | >16 | N.D. | 16 | N.D. | <0.31 (Syn) |
| *P. aeruginosa* ATCC BAA 2108 | N.D. | N.D. | N.D. | N.D. | N.D. | N.D. |
| *P. aeruginosa* ATCC 27853 | N.D. | N.D. | N.D. | N.D. | N.D. | N.D. |

*N.D. refers to "not determined"

**Table S3.** Differentially expressed genes (DEGs) of *E. coli* ATCC BAA 2471 upon treatment of D-PP (8 µg/mL) vs untreated control with a $Log_2$(Fold change) ≥ 1 and $Log_2$(Fold change) ≤ -1 and P value <0.05.

| DEGs | Gene Product | log2(Fold change) |
|---|---|---|
| aceK | isocitrate dehydrogenase kinase/phosphatase | -1.04674 |
| prpB | 2-methylisocitrate lyase | 1.136358 |
| prpC | 2-methylcitrate synthase | 1.254795 |
| prpD | 2-methylcitrate dehydratase | 1.246424 |
| acs_2 | Unknown | 1.448194 |
| yaiY | DUF2755 domain-containing inner membrane protein YaiY | 1.96911 |
| glnK | uridylyl-[protein PII-2] // nitrogen regulatory protein PII-2 | 1.102003 |
| ybhF | ABC exporter ATP binding subunit YbhF | 1.026614 |
| cecR | DNA-binding transcriptional dual regulator CecR | 2.145607 |
| rmf | ribosome modulation factor | 1.195957 |
| ompX_2 | Protein-binding (outer membrane protein X) ompX | 2.357283 |
| bdm | biofilm-dependent modulation protein | 1.816852 |
| mtlK | Unknown | 1.085695 |
| spaP | Unknown | -1.17568 |
| mxiC | Unknown | -1.10263 |
| glgS | surface composition regulator | 1.22459 |
| mtr | tryptophan:H<sup>+</sup> symporter Mtr | -1.11622 |
| dctR | putative DNA-binding transcriptional regulator DctR | -1.02056 |
| asnA | asparagine synthetase A | 1.207694 |

**Table S4.** Differentially expressed genes (DEGs) of *E. coli* ATCC BAA 2471 upon treatment of D-PP (16µg/mL) vs untreated control with a $Log_2$(Fold change) ≥ 1 and $Log_2$(Fold change) ≤ -1 and P value <0.05.

| DEGs | Gene Product | log2(Fold change) |
|---|---|---|
| gldA | L-1,2-propanediol dehydrogenase / glycerol dehydrogenase | 1.339436 |
| fryA_1 | putative PTS multiphosphoryl transfer protein FryA | 1.106899 |
| aceB | malate synthase A | -1.43464 |
| aceA | isocitrate lyase | -1.85769 |
| aceK | isocitrate dehydrogenase kinase/phosphatase | -2.08356 |
| soxS | DNA-binding transcriptional dual regulator SoxS | 1.287077 |
| soxR | DNA-binding transcriptional dual regulator SoxR | 1.120543 |

| | | |
|---|---|---|
| phnK | ATP-binding cassette protein PhnK | 1.232363 |
| fumB | fumarase B | 1.200309 |
| dcuB | anaerobic C4-dicarboxylate transporter DcuB | 1.036375 |
| frdB | fumarate reductase iron-sulfur protein | 1.091474 |
| treC | trehalose-6-phosphate hydrolase | -1.25013 |
| treB | trehalose-specific PTS enzyme IIBC component | -1.23421 |
| rraB | ribonuclease E inhibitor protein B | 1.419809 |
| fecA | ferric citrate outer membrane transporter | -1.018 |
| lgoR | putative DNA-binding transcriptional regulator LgoR | 1.036473 |
| yjjW | putative glycyl-radical enzyme activating enzyme YjjW | 1.141382 |
| araD | L-ribulose-5-phosphate 4-epimerase AraD | 1.05051 |
| ivy | periplasmic chaperone, inhibitor of vertebrate C-type lysozyme | 1.066083 |
| prpB | 2-methylisocitrate lyase | 1.482858 |
| prpC | 2-methylcitrate synthase | 1.434503 |
| prpD | 2-methylcitrate dehydratase | 1.584821 |
| acs_2 | acetyl-CoA synthetase (AMP-forming) | 1.732089 |
| yaiY | DUF2755 domain-containing inner membrane protein YaiY | 2.471892 |
| decR | DNA-binding transcriptional activator DecR | 1.269846 |
| glnK | uridylyl-[protein PII-2] // nitrogen regulatory protein PII-2 | 2.782331 |
| amtB | ammonium transporter | 1.708328 |
| fepA | ferric enterobactin outer membrane transporter | -1.34061 |
| fepE | polysaccharide co-polymerase family protein FepE | -1.19198 |
| fepC | ferric enterobactin ABC transporter ATP binding subunit | -1.35378 |
| uspG | universal stress protein G | 1.299706 |
| glnQ_1 | L-glutamine ABC transporter ATP binding subunit | -1.00788 |
| ybgD_1 | putative fimbrial protein YbgD | -1.34274 |
| bioC | malonyl-acyl carrier protein methyltransferase | -1.02513 |
| ybhS | ABC exporter membrane subunit YbhS | 1.097324 |
| ybhF | ABC exporter ATP binding subunit YbhF | 1.470927 |
| cecR | DNA-binding transcriptional dual regulator CecR | 2.72843 |
| opgE | phosphoethanolamine transferase OpgE | -1.15724 |
| nfsA | NADPH-dependent nitro/quinone reductase NfsA | 1.847858 |
| rimK | ribosomal protein S6 modification protein | 1.242624 |
| dmsA | dimethyl sulfoxide reductase subunit A | 1.023293 |
| dmsB_1 | dimethyl sulfoxide reductase subunit B | 1.072569 |
| ompF | outer membrane porin F | -1.59581 |
| rmf | ribosome modulation factor | 1.042651 |
| gnsA | putative phosphatidylethanolamine synthesis regulator GnsA | -1.09466 |
| csgF | curli assembly component CsgF | -1.61985 |
| csgE | curli assembly component CsgE | -1.38342 |
| ompX_2 | Protein-binding (outer membrane protein X) ompX | 1.805383 |
| ydfO | Qin prophage; DUF1398 domain-containing protein YdfO | -1.42366 |
| prmC | protein-(glutamine-N5) methyltransferase | -1.02544 |
| adhE | aldehyde/alcohol dehydrogenase AdhE | 1.089288 |
| ompW | outer membrane protein W | 1.348583 |

| | | |
|---|---|---|
| puuC | γ-glutamyl-γ-aminobutyraldehyde dehydrogenase | -1.06715 |
| uspF | Universal stress protein F | 1.140372 |
| nifJ | Unknown | 1.036063 |
| bdm | biofilm-dependent modulation protein | 2.27612 |
| fimH | type 1 fimbriae D-mannose specific adhesin | -1.07452 |
| marR | DNA-binding transcriptional repressor MarR | 1.4316 |
| marA | DNA-binding transcriptional dual regulator MarA | 1.269657 |
| fumC | fumarase C | 1.983895 |
| cnu | H-NS- and StpA-binding protein | -1.09777 |
| uvrC | UvrABC excision nuclease subunit C | -1.09456 |
| yoeB | ribosome-dependent mRNA interferase toxin YoeB | 1.34653 |
| mdtD | putative multidrug efflux pump MdtD | 1.100727 |
| preT | dihydropyrimidine dehydrogenase (NAD+) subunit PreT | 2.000927 |
| preA | dihydropyrimidine dehydrogenase (NAD+) subunit PreA | 2.032745 |
| mtlK | Unknown | 1.641019 |
| inaA | putative lipopolysaccharide kinase InaA | 1.27971 |
| glpA | anaerobic glycerol-3-phosphate dehydrogenase subunit A | 1.309541 |
| evgS | sensor histidine kinase EvgS | -1.15445 |
| hscB | [Fe-S] cluster biosynthesis co-chaperone HscB | -1.25506 |
| iscA | iron-sulfur cluster insertion protein IscA | -1.00911 |
| grcA | stress-induced alternate pyruvate formate-lyase subunit | 1.534379 |
| nrdF | ribonucleoside-diphosphate reductase 2 subunit β | -1.31156 |
| malY_2 | negative regulator of MalT activity/cystathionine and beta; -lyase | -1.08501 |
| malX_2 | PTS enzyme IIBC component MalX | -1.00865 |
| argA | N-acetylglutamate synthase | 1.283747 |
| spaP | Unknown | -1.2531 |
| invA | Unknown | -1.12332 |
| ygeX | 2,3-diaminopropionate ammonia-lyase | 1.20763 |
| argE_2 | acetylornithine deacetylase | 1.498595 |
| ygfK | putative oxidoreductase, Fe-S subunit | 1.190816 |
| ssnA | putative aminohydrolase SsnA | 1.265057 |
| idi | isopentenyl-diphosphate Δ-isomerase | 2.01309 |
| ansB | L-asparaginase 2 | 1.511557 |
| yqhD | NADPH-dependent aldehyde reductase YqhD | 1.286171 |
| glgS | surface composition regulator | 2.297727 |
| lldR_1 | DNA-binding transcriptional dual regulator LldR | 1.225849 |
| tdcE | 2-oxobutanoate formate-lyase/pyruvate formate-lyase 4 | 1.036674 |
| tdcD | propionate kinase | 1.114026 |
| tdcC | threonine/serine:H+ symporter TdcC | 1.248024 |
| tdcB | catabolic threonine dehydratase | 1.385055 |
| tsaR_2 | Unknown | 1.248297 |
| mtr | tryptophan:H+ symporter Mtr | -1.41958 |
| bhsA_3 | DUF1471 domain-containing multiple stress resistance outer membrane protein BhsA | 1.325885 |
| yhdW | putative ABC transporter periplasmic binding protein YhdW | -1.09096 |

| | | |
|---|---|---|
| glpD | aerobic glycerol 3-phosphate dehydrogenase | 1.01715 |
| gntR | DNA-binding transcriptional repressor GntR | 1.08458 |
| zntA | Zn2+/Cd2+/Pb2+ exporting P-type ATPase | 1.003429 |
| nikB | nickel ABC transporter membrane subunit NikB | 1.257646 |
| nikC | nickel ABC transporter membrane subunit NikC | 1.256835 |
| rbbA | ribosome-associated ATPase | 1.199561 |
| dctR | putative DNA-binding transcriptional regulator DctR | -2.05623 |
| sapB_2 | putrescine ABC exporter membrane subunit SapB | -1.71156 |
| gadW | DNA-binding transcriptional dual regulator GadW | -1.22194 |
| dppD | dipeptide ABC transporter ATP binding subunit DppD | -1.02625 |
| dppC | dipeptide ABC transporter membrane subunit DppC | -1.31945 |
| dppB | dipeptide ABC transporter membrane subunit DppB | -1.35375 |
| hokA_2 | small toxic polypeptide | -1.54639 |
| mtlR | transcriptional repressor MtlR | 1.675107 |
| ibpB | small heat shock protein IbpB | 1.549134 |
| asnA | asparagine synthetase A | 1.646912 |
| chuR_2 | Unknown | 1.144344 |
| glnA | adenylyl-[glutamine synthetase] / glutamine synthetase | 1.424684 |
| ysdC | Unknown | 1.362384 |
| manP_2 | Unknown | 1.139873 |
| tetA | Unknown | 1.530235 |
| tetR | Unknown | 1.39728 |
| mntB_1 | Unknown | -1.07804 |
| hin_1 | Unknown | 1.04418 |

**Table S5.** Differentially expressed genes (DEGs) of *K. pneumoniae* ATCC BAA 2473 upon treatment of D-PP (8 μg/mL) vs untreated control with a $Log_2$(Fold change) ≥ 1 and $Log_2$(Fold change) ≤ -1 and P value <0.05.

| DEGs | Gene Product | log2(Fold change) |
|---|---|---|
| ygdR | Unknown | 1.95163 |
| malK | maltose/maltodextrin ABC transporter ATP-binding protein MalK | 1.90334 |
| raiA | ribosome-associated translation inhibitor RaiA | 1.74300 |
| ygdR | Unknown | 1.63580 |
| ycaC | Unknown | 1.57531 |
| cadA | Unknown | 1.55233 |
| malE | maltose/maltodextrin ABC transporter substrate-binding protein MalE | 1.49195 |
| yebV | Unknown | 1.48283 |
| ibpB | heat shock chaperone IbpB | 1.40114 |
| ycgR | Unknown | 1.36543 |
| lamB | porin LamB | 1.29394 |
| higB2 | Unknown | 1.28162 |
| cadB | lysine decarboxylase CadA | 1.27380 |

| | | |
|---|---|---|
| nemR | Unknown | 1.23995 |
| lpp | Unknown | 1.23715 |
| maa | maltose O-acetyltransferase | 1.23019 |
| intA | Unknown | 1.19504 |
| dinI | DNA damage-inducible protein I | 1.16194 |
| asnB | asparagine synthase B | 1.12911 |
| hha | hemolysin expression modulator Hha | 1.12858 |
| glgS | cell surface composition regulator GlgS | 1.11175 |
| pspG | envelope stress response protein PspG | 1.10854 |
| mngR | Unknown | 1.08986 |
| cusC | Unknown | 1.08724 |
| bhsA | Unknown | 1.07407 |
| yaiY | Unknown | 1.06497 |
| gcvA | transcriptional regulator GcvA | 1.06351 |
| malG | maltose ABC transporter permease MalG | 1.05898 |
| ycgZ | Unknown | 1.04629 |
| ttdT | Unknown | 1.04580 |
| yraI | Unknown | 1.04120 |
| nirD | nitrite reductase small subunit NirD | 1.03436 |
| malX | Unknown | 1.03059 |
| soxS | superoxide response transcriptional regulator SoxS | 1.03022 |
| alsD | Unknown | 1.02939 |
| chbA | PTS *N,N'*-diacetylchitobiose transporter subunit IIA | 1.02315 |
| ibpA | heat shock chaperone IbpA | 1.01500 |
| osmB | osmotically-inducible lipoprotein OsmB | 1.00422 |
| ugpA | sn-glycerol-3-phosphate ABC transporter permease UgpA | -1.00195 |
| astB | N-succinylarginine dihydrolase | -1.00306 |
| pduC | propanediol dehydratase large subunit PduC | -1.00366 |
| pcaB | Unknown | -1.00460 |
| astA | arginine N-succinyltransferase | -1.00881 |
| ahr | Unknown | -1.00897 |
| ycjG | L-Ala-D/L-Glu epimerase | -1.01064 |
| traI | conjugative transfer relaxase/helicase TraI | -1.02349 |
| uao | Unknown | -1.02490 |
| fucI | L-fucose isomerase | -1.03424 |
| entH | proofreading thioesterase EntH | -1.04308 |
| cysG | Unknown | -1.04781 |
| ghrB | glyoxylate/hydroxypyruvate reductase GhrB | -1.05168 |
| astD | succinylglutamate-semialdehyde dehydrogenase | -1.05172 |
| fumC | class II fumarate hydratase | -1.05434 |
| clpC | Unknown | -1.05445 |
| rbsA | ribose ABC transporter ATP-binding protein RbsA | -1.05503 |
| selB | selenocysteine-specific translation elongation factor | -1.05893 |
| gbuA | Unknown | -1.06296 |
| soxC | Unknown | -1.06907 |

| | | |
|---|---|---|
| yceM | Unknown | -1.06945 |
| feaB | phenylacetaldehyde dehydrogenase | -1.07159 |
| tauB | taurine ABC transporter ATP-binding subunit | -1.07423 |
| dipZ | Unknown | -1.07636 |
| amnD | Unknown | -1.09922 |
| murG | undecaprenyldiphospho-muramoylpentapeptide beta-N-acetylglucosaminyltransferase | -1.09982 |
| hyuA | Unknown | -1.10201 |
| ugpC | sn-glycerol-3-phosphate ABC transporter ATP-binding protein UgpC | -1.11131 |
| hmuU | hemin ABC transporter membrane protein HmuU | -1.11246 |
| eutN | ethanolamine utilization microcompartment protein EutN | -1.11427 |
| kgtP | Unknown | -1.11921 |
| ddrA | Unknown | -1.12018 |
| noc | Unknown | -1.12547 |
| tdcD | propionate kinase | -1.12557 |
| sucC | ADP-forming succinate--CoA ligase subunit beta | -1.12580 |
| ddpC | Unknown | -1.12810 |
| dppC | dipeptide ABC transporter permease DppC | -1.12997 |
| phnG | phosphonate C-P lyase system protein PhnG | -1.13017 |
| nupC | nucleoside permease NupC | -1.13770 |
| aceA | isocitrate lyase | -1.13943 |
| pcaF | 3-oxoadipyl-CoA thiolase | -1.14256 |
| pduE | propanediol dehydratase small subunit PduE | -1.14315 |
| mdh | malate dehydrogenase | -1.14338 |
| cobT | nicotinate-nucleotide--dimethylbenzimidazole phosphoribosyltransferase | -1.14373 |
| ybgJ | Unknown | -1.15980 |
| cbiE | Unknown | -1.16583 |
| menC | o-succinylbenzoate synthase | -1.17926 |
| lvr | Unknown | -1.19656 |
| cobT | nicotinate-nucleotide--dimethylbenzimidazole phosphoribosyltransferase | -1.19927 |
| proY | proline-specific permease ProY | -1.20171 |
| menE | o-succinylbenzoate--CoA ligase | -1.20584 |
| cbiL | Unknown | -1.22552 |
| pduD | propanediol dehydratase medium subunit PduD | -1.22708 |
| ccmH | Unknown | -1.24582 |
| yehX | Unknown | -1.25008 |
| bcsC | cellulose biosynthesis protein BcsC | -1.25070 |
| astD | succinylglutamate-semialdehyde dehydrogenase | -1.25299 |
| bdcA | SDR family oxidoreductase | -1.25922 |
| dadA | Unknown | -1.27167 |
| ccmC | Unknown | -1.27167 |
| menI | 1,4-dihydroxy-2-naphthoyl-CoA hydrolase | -1.29444 |
| feaB | phenylacetaldehyde dehydrogenase | -1.31581 |

| | | |
|---|---|---|
| pduA | propanediol utilization microcompartment protein PduA | -1.35155 |
| ygfF | Unknown | -1.36590 |
| pduB | propanediol utilization microcompartment protein PduB | -1.37711 |
| entF | apo-serine activating enzyme EntF | -1.37882 |
| sacA | Unknown | -1.38297 |
| traQ | type-F conjugative transfer system pilin chaperone TraQ | -1.40349 |
| amaB | Unknown | -1.43998 |
| traI | conjugative transfer relaxase/helicase TraI | -1.46351 |
| oadB | Unknown | -1.46405 |
| pduA | propanediol utilization microcompartment protein PduA | -1.53428 |
| potA | spermidine/putrescine ABC transporter ATP-binding protein PotA | -1.54383 |
| yxeP | Unknown | -1.59587 |
| proA | glutamate-5-semialdehyde dehydrogenase | -1.67156 |
| recD | exodeoxyribonuclease V subunit alpha | -1.76413 |
| ampG | muropeptide MFS transporter AmpG | -1.99864 |
| mbtI | Unknown | -2.02903 |
| can | carbonate dehydratase | -2.03339 |
| ais | Unknown | -2.03477 |
| pikAV | Unknown | -2.14384 |
| dhbE | Unknown | -2.24633 |

**Table S6.** Differentially expressed genes (DEGs) of *K. pneumoniae* ATCC BAA 2473 upon treatment of D-PP (16 µg/mL) vs untreated control with a $Log_2$(Fold change) ≥ 1 and $Log_2$(Fold change) ≤ -1 and P value <0.05.

| DEGs | Gene Product | Log2(Fold change) |
|---|---|---|
| malP | maltodextrin phosphorylase | 1.03734 |
| aglB | Unknown | 1.45846 |
| dppA | Unknown | -1.02149 |
| dppB | dipeptide ABC transporter permease DppB | -1.44904 |
| dppC | dipeptide ABC transporter permease DppC | -1.67497 |
| dppD | dipeptide ABC transporter ATP-binding protein | -1.16006 |
| puuE | allantoinase PuuE | -1.40278 |
| ugpC | sn-glycerol-3-phosphate ABC transporter ATP-binding protein UgpC | -1.21320 |
| nirD | nitrite reductase small subunit NirD | 1.06646 |
| lsrA | autoinducer 2 ABC transporter ATP-binding protein LsrA | -1.01196 |
| lsrC | autoinducer 2 ABC transporter permease LsrC | -1.28325 |
| lsrD | Unknown | -1.47121 |
| lsrB | autoinducer 2 ABC transporter substrate-binding protein LsrB | -1.41625 |
| lsrF | 3-hydroxy-5-phosphonooxypentane-2,4-dione thiolase | -1.11838 |
| glgS | cell surface composition regulator GlgS | 1.22597 |
| ycgR | Unknown | 1.02878 |

| | | |
|---|---|---|
| sacA | Unknown | -1.21648 |
| recD | exodeoxyribonuclease V subunit alpha | -1.05046 |
| pduU | propanediol utilization microcompartment protein PduU | 1.29288 |
| rsxC | putative ion-translocating oxidoreductase complex subunit RsxC | 1.19089 |
| pduL | Unknown | 1.03218 |
| pduA | propanediol utilization microcompartment protein PduA | 1.49728 |
| ddrA | Unknown | 1.90952 |
| pduE | propanediol dehydratase small subunit PduE | 1.90063 |
| pduD | propanediol dehydratase medium subunit PduD | 2.13794 |
| pduC | propanediol dehydratase large subunit PduC | 1.60559 |
| pduB | propanediol utilization microcompartment protein PduB | 1.89471 |
| pduA | propanediol utilization microcompartment protein PduA | 2.02882 |
| pduF | propanediol diffusion facilitator PduF | 1.54518 |
| soxC | Unknown | -1.00346 |
| raiA | ribosome-associated translation inhibitor RaiA | 1.00199 |
| dinI | DNA damage-inducible protein I | 1.04612 |
| intA | Unknown | 1.03975 |
| mlaA | phospholipid-binding lipoprotein MlaA | 1.08928 |
| cadA | Unknown | 1.22849 |
| tetA | Unknown | 1.08580 |
| cimH | Unknown | 1.77253 |
| ygdR | Unknown | 1.07436 |
| rbsA | ribose ABC transporter ATP-binding protein RbsA | -1.17574 |
| pspG | envelope stress response protein PspG | 1.01536 |
| aceK | bifunctional isocitrate dehydrogenase kinase/phosphatase | -1.33908 |
| aceA | isocitrate lyase | -1.62410 |
| aceB | malate synthase A | -1.29882 |
| asnA | aspartate--ammonia ligase | 1.26565 |
| ybdO | Unknown | 1.27351 |
| asnB | asparagine synthase B | 1.03670 |
| sucC | ADP-forming succinate--CoA ligase subunit beta | -1.02259 |
| proY | proline-specific permease ProY | -1.15348 |
| cecR | transcriptional regulator CecR | 1.18064 |
| bhsA | Unknown | 1.20971 |
| feaB | phenylacetaldehyde dehydrogenase | -1.16253 |
| fnr | fumarate/nitrate reduction transcriptional regulator Fnr | 1.00418 |
| pcaI | Unknown | -1.06739 |
| pcaJ | Unknown | -1.07416 |
| mngR | Unknown | 1.11785 |
| narW | nitrate reductase molybdenum cofactor assembly chaperone | -1.03011 |
| nemR | Unknown | 1.08977 |
| ccmH | Unknown | -1.04547 |
| dadA | Unknown | -1.58162 |

| | | |
|---|---|---|
| amnD | Unknown | -1.53540 |
| ais | Unknown | -2.26128 |
| kgtP | Unknown | -1.45338 |
| livH | high-affinity branched-chain amino acid ABC transporter permease LivH | -1.04731 |
| mglC | galactose/methyl galactoside ABC transporter permease MglC | -1.03993 |
| mglA | galactose/methyl galactoside ABC transporter ATP-binding protein MglA | -1.04209 |
